# Blood-brain barrier dysfunction in aging induces hyper-activation of TGF-beta signaling and chronic yet reversible neural dysfunction

**DOI:** 10.1101/537431

**Authors:** V.V. Senatorov, A.R. Friedman, D.Z. Milikovsky, J. Ofer, R. Saar-Ashkenazy, A. Charbash, N. Jahan, G. Chin, E. Mihaly, J.M. Lin, H.J. Ramsay, A. Moghbel, M.K. Preininger, C.R. Eddings, H.V. Harrison, R. Patel, Y. Shen, H. Ghanim, H. Sheng, R. Veksler, P.H. Sudmant, A. Becker, B. Hart, M.A. Rogawski, A. Dillin, A. Friedman, D. Kaufer

## Abstract

Aging involves a decline in neural function that contributes to cognitive impairment and disease. However, the mechanisms underlying the transition from a young-and-healthy to aged-and-dysfunctional brain are not well understood. Here, we report breakdown of the vascular blood-brain barrier (BBB) in aging humans and rodents, which begins as early as middle age and progresses to the end of the lifespan. Gain-of-function and loss-of-function manipulations show that this BBB dysfunction triggers hyperactivation of transforming growth factor β (TGFβ) signaling in astrocytes, which is necessary and sufficient to cause neural dysfunction and age-related pathology. Specifically, infusion of the serum protein albumin into the young brain (mimicking BBB leakiness) induced astrocytic TGFβ signaling and an aged brain phenotype including aberrant electrocorticographic activity, vulnerability to seizures, and cognitive impairment. Furthermore, conditional genetic knockdown of astrocytic TGFβ receptors, or pharmacological inhibition of TGFβ signaling, reversed these symptomatic outcomes in aged mice. Finally, we found that this same signaling pathway is activated in aging human subjects with BBB dysfunction. Our study identifies dysfunction in the neurovascular unit as one of the earliest triggers of neurological aging, and demonstrates that the aging brain may retain considerable latent capacity which can be revitalized by therapeutic inhibition of TGFβ signaling.

## Introduction

Aging involves cognitive deterioration which poses an increasing healthcare burden on today’s society with its prolonged life expectancy. Despite calls for a better understanding of brain aging and new therapeutic targets, the underlying mechanisms that cause decline in neural function in aging are still not understood. We approached this topic by exploring the causal involvement of the blood-brain barrier (BBB) pathology in age-related brain dysfunction. The BBB is a tightly-regulated interface composed of specialized endothelial cells, pericytes and astrocytic end-feet that form a protective sheath around brain capillaries. By restricting the free diffusion of blood-borne molecules, the BBB establishes a sequestered brain microenvironment, including the precisely balanced ionic concentrations needed for neural activity, the compartmentalization of brain-specific growth factors and signaling molecules, and the immune-privileged brain environment *(1, 2)*. Thus, the BBB is a fundamental and essential component of healthy brain function.

Alarming observations of widespread BBB breakdown in aging patients were first reported in the 1970s *(3)*, raising the possibility that vascular leakiness and infiltration of toxic blood-borne molecules into the brain could cause neural impairments and contribute to diseases such as dementia *(2, 4–7)*. This theory has been primarily supported by human clinical evidence showing that BBB breakdown in aging individuals is strongly correlated with cognitive decline and Alzheimer’s disease *(7–12)*. Furthermore, this association has been localized to functional brain sub-regions: BBB breakdown specifically in the hippocampus is associated with significant mild cognitive impairment *(13)* – suggesting that BBB dysfunction could be a cause of impairment within impacted tissues. However, these human studies have also yielded contradictory results, are complicated by variability in population and study methodology *(14, 15)*, and are inherently correlative. There have been very few rodent studies that assess BBB status in aging *(16)*, and to our knowledge, no studies that test mechanistic hypotheses about how BBB breakdown affects brain function in aging. A mechanistic understanding of the biological consequences of BBB breakdown is critical to discern whether the association between BBB dysfunction and cognitive decline is spurious, correlative, or causal. Furthermore, mechanistic insights would have the potential to reveal new druggable targets for the intractable health problems of age-related cognitive decline and dementia. Indeed, an improved understanding of vascular contributions to cognitive decline and dementia has been designated as one of the highest priority research areas for new breakthroughs in Alzheimer’s disease and related dementias *(17)*.

While little is known about the consequences of BBB breakdown in aging, there are several other disease contexts that involve BBB dysfunction, which could reveal relevant candidate mechanisms. In particular, traumatic brain injury (TBI) not only causes severe BBB breakdown *(18–20)*, but also involves secondary symptoms of cognitive impairment and increases risk for dementia. In rodent models of TBI and leaky BBB, blood-borne proteins that infiltrate into the brain cause a robust injury response by activating the transforming growth factor beta (TGFβ) signaling pathway. Key mediators of this response include the serum protein fibrinogen, which carries and releases latent TGFβ in the brain *(21)*, and albumin, which binds to the TGFβ receptor and activates signaling *(22)*. In both cases, astrocytes act as the primary responders that detect the blood-borne ligands and transduce TGFβ signaling *(21–23)*. In turn, activated astrocytes release inflammatory cytokines and more TGFβ1 *(22, 24, 25)*, form glial scars *(21)*, and remodel neural circuits to cause hyperexcitability and dysfunction *(22, 24, 26–28)*. These studies point to the TGFβ signaling pathway as a candidate mechanism that could play a role in pathological outcomes following BBB breakdown in aging. To test this hypothesis, we used genetic and pharmaceutical interventions to test the causality of BBB dysfunction and TGFβ signaling in progressive neural dysfunction across the lifespan of naturally aging mice.

## Results

### Progressive BBB dysfunction and albumin extravasation in the hippocampus starts in middle age

To establish the time course of age-related BBB decline, we quantified extravasation of serum albumin, an established marker of BBB permeability *(29–31)* which also plays a mechanistic role in triggering TGFβ signaling *(22)*. We focused our analysis on the hippocampus – a key brain region associated with age-related memory decline *(13)*. Albumin was effectively absent from the hippocampus of young mice, but was first detectable in the aging hippocampus starting as early as 12 months (“middle age”) and consistently elevated in aging up to two years, near the end of the lifespan (Fig. 1A and Fig. S1A). These results indicate that BBB dysfunction appears earlier than has been appreciated, placing it among the earliest known biomarkers of aging in the rodent brain. The onset of BBB dysfunction at around 12 months corresponds with other early biological hallmarks of aging, including the typical onset of reproductive senescence in female mice, as well as the earliest appearance of mild cognitive impairments in various behavior tasks *(32–37)*. To confirm age-related BBB dysfunction, we raised a separate cohort of aged mice and used an alternate method to quantify BBB permeability, based on detecting leakage of a fluorescent tracer, Evans blue (EB), into the brain after i.v. injection *(38)*. Compared to 2-4 month old young mice, 12 and 21-24 month old aged mice showed significantly elevated levels of EB leakage into the brain (Fig. S1B-C), thus reproducing our finding of age-related BBB breakdown by an independent method.

**Fig. 1.**
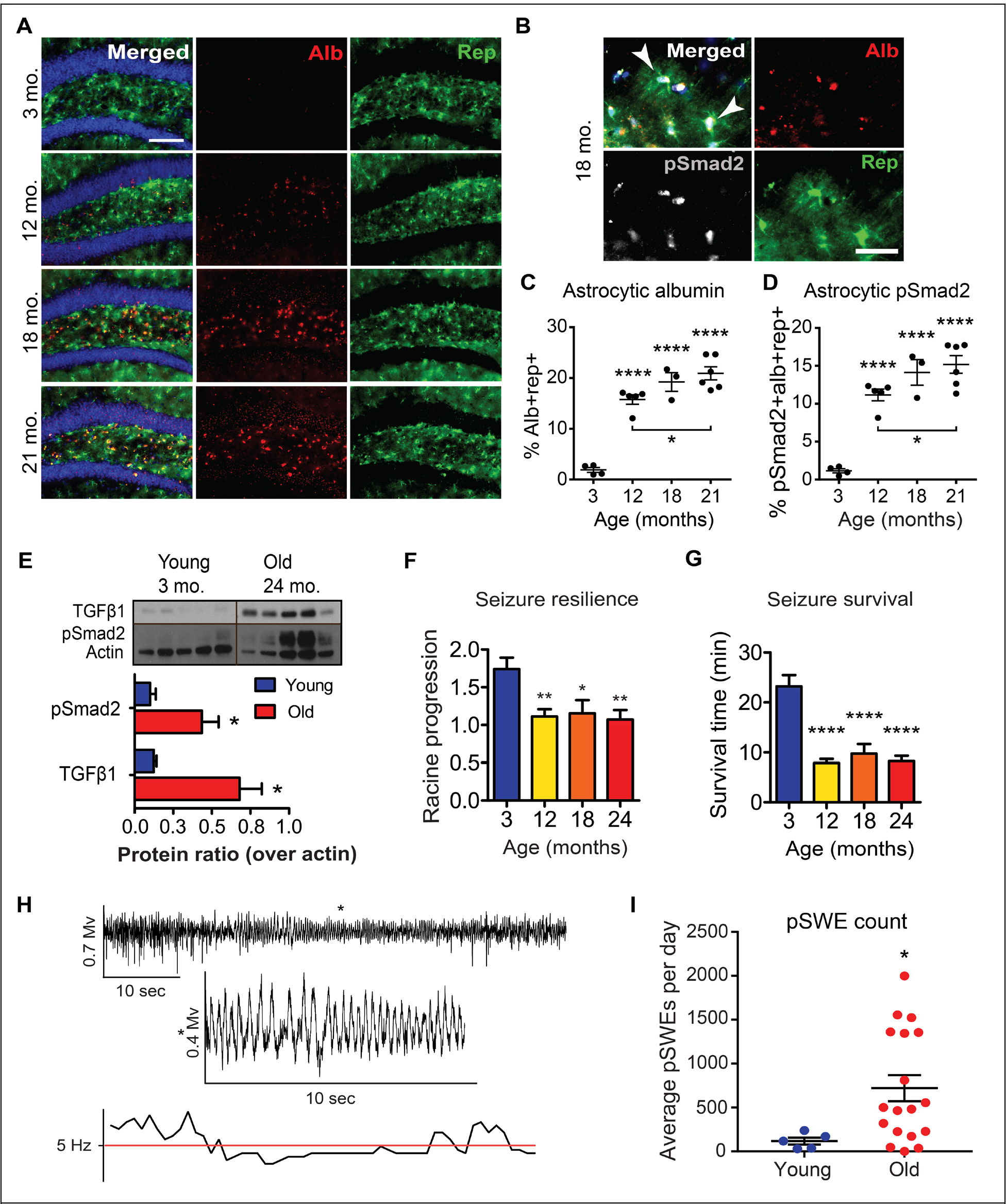
Progressive BBB dysfunction in aging mice is associated with elevated TGFβ signaling and aberrant network activity. (A) Immunofluorescent staining used to quantify progression of BBB decline in the aging mouse hippocampus at 3, 12, 18, and 21 months old. Astrocytes are labeled by GFP expressed transgenically under the pan-astrocytic promoter Aldh1L1 (Rep-aldh1L1-eGFP), albumin is labeled by immunostaining, and cell nuclei by DAPI (blue). Scale bar = 100 μm. (B) Immunostaining was used to investigate co-localization of albumin and pSmad2 (arrows) in Rep-aldh1L1-eGFP astrocytes (Scale bar = 30 μm). (C) Albumin localized in astrocytes increases with age (ANOVA, p < 0.0001). (D) Aging mice show an increase in the level of pSmad2 colabeled with albumin in astrocytes (ANOVA, p = <0.0001). For all measures, groups were compared by Bonferroni post-hoc test. Sample sizes are n = 4 (3 mo); 5 (12 mo); 3 (18 mo); and 6 (21 mo). (E) Western blot analyzing TGFβ1 and pSmad2, outputs of the TGFβ signaling pathway, in hippocampus from young and old mice. Western blot densitometry shows that pSmad2 (t-test, p=0.019) and TGFβ (t-test with Welch’s correction, p = 0.018) protein levels were elevated in the aged mouse hippocampus (n = 5). (F) At 3, 12, 18, and 24 months seizures were induced in mice by PTZ injection to assay hyperexcitability. Seizure severity in mice was quantified by fitting linear regression slopes to measures of progression through each stage of seizure in the modified Racine scale. Aged mice (12-24 months) had decreased linear regression slopes indicating faster progression through all stages of seizure severity (1-way ANOVA, p = 0.0011 with Bonferroni posttest; n = 13 (3 mo), 10 (12 mo), 8 (18 mo and 24 mo)). (G) Latency to mortality caused by severe seizures was significantly faster in 12-24 month old groups compared to young mice (1-way ANOVA, p < 0.0001 with Bonferroni posttest). (H) A representative trace shows a pSWE with slow-wave activity less than 5 Hz within a 10 second window (marked with *). (I) Aged mice showed a significantly elevated number of pSWEs compared to young mice. For all tests, *p<0.05, **p<0.01, ***p<0.005, ****p<0.001.

### Age-dependent albumin accumulation is cell-type specific

Albumin is known to accumulate in astrocytes when it extravasates through a dysfunctional BBB *(22, 24, 39)*. We therefore used a transgenic mouse line that comprehensively labels astrocytes via the Aldh1L1 (Aldehyde Dehydrogenase 1 Family Member L1) promoter (Rep-Aldh1L1) *(40, 41)*. Expression of this marker was stable and did not change across the lifespan (Fig. S1D). To estimate the specificity of albumin uptake in different cell types, we performed immunostaining for markers of microglia (Iba1), oligodendrocytes (CAII), and neurons (NeuN), and quantified the percent of all albumin+ cells for each cell type in the aged (18-24 months) mouse hippocampus. Albumin co-localized predominantly with Aldh1L1-positive astrocytes, accounting for approximately 60% of albumin-labeled cells, while each of the other cell types co-labeled approximately 20% of all albumin-positive cells (Fig. S1E). Concurrent with the time course of overall albumin extravasation and BBB dysfunction, significant astrocytic uptake of albumin was first detected at 12 months, and further increased with age up to 21 months old (Fig. 1C).

### Age-dependent activation of aberrant TGFβ signaling in astrocytes

Albumin endocytosis into astrocytes is mediated by binding to the TGFβ receptor II (TGFβR) subunit, which in turn activates the TGFβRI ALK5 *(22, 28)* to induce phosphorylation of Smad2 (pSmad2) and carry out signal transduction of the ALK5-TGFβ signaling cascade. Further, this activation also increases the production of TGFβ1 in astrocytes (Weissberg et al., 2015) and activation of latent TGFβ1 protein from extra-cellular matrix *(26)*, yielding an increase in the canonical ligand of TGFβR and therefore amplification of the TGFβ cascade. Hence, we next investigated the relationship between albumin uptake and TGFβ signaling in the aged mouse brain by quantifying immunolabeled phosphorylated Smad2 protein (pSmad2). Concurrent with the time course of albumin extravasation, aging mice showed a progressive increase in the levels of pSmad2 co-localized with albumin-positive astrocytes (Fig. 1B and D). Activation of the TGFβ pathway was further quantified by Western blot, showing increased levels of pSmad2 in the hippocampus of old mice, compared to young, and increased levels of active TGFβ1, a positive feedback output of the TGFβ pathway (Fig. 1E).

### Network hyperexcitability in aged mice

Based on the finding that BBB dysfunction and TGFβ signaling causes hyperexcitability after head injury *(22, 24, 26–28, 39)*, we hypothesized that similar hyperexcitability may be triggered by BBB decline and contribute to cognitive impairment in aging mice. Indeed, hippocampal hyperexcitability is an early biomarker of mild cognitive impairment in humans that precedes progression to AD *(42, 43)*, and is also an early marker of disease progression in rodent AD models *(44, 45)*. Thus, we used the pentylenetetrazol (PTZ) seizure assay to investigate the time course for onset and progression of hyperexcitability in aging mice. In this assay, injection of PTZ, a non-competitive GABA receptor blocker, induces seizures that have been shown to be a reliable readout of underlying hippocampal hyperexcitability associated with aging and AD models *(45)* (i.e. mice with higher hyperexcitability exhibit lower induced seizure threshold). Compared to 3 month old mice, aged mice showed increased severity of induced seizures (quantified based on rate of progression through the Racine seizure severity scale, Fig. S1F), beginning at the 12-month-old time point (Fig. 1F). Furthermore, old mice were highly vulnerable to mortality from severe seizures, with a significantly shorter latency to mortality (Fig. 1G).

### Aberrant paroxysmal slow wave events in aged mice

Next, we sought to measure and characterize hyperexcitability directly via electrophysiology. To do so, we recorded telemetric electrocorticography (ECoG) using epidural electrodes implanted in young (3 months old) and old (18-24 months old) mice over a period of 5 days in the home cage. We found that aged mice showed an increase in the relative power of slow wave activity (<5 Hz) (Fig. S1G), similar to EEG slowing described in human dementia patients *(46–48)*, which is thought to reflect dysfunctional neural networks. Detailed analysis of this aberrant ECoG signal revealed that the slow-wave activity was not continuous, but rather manifested in discrete, transient paroxysmal slow wave episodes (pSWEs, median frequency < 5 Hz; Fig. 1H), which were significantly elevated in aged mice relative to young (Fig. 1I). These pSWEs, characterized by a median power of less than 5 Hz over 10 consecutive seconds, were similar to paroxysmal ECoG events that have been observed in epileptogenic animals that have hippocampal hyperexcitability *(49)*. Given the potential association between hyperexcitability and seizures, we also used an automated seizure detection algorithm *(24, 49)* to search for spontaneous seizures in the ECoG recordings, and found that aged mice had no seizure events and were not epileptic. Thus, the pSWEs constitute distinct subclinical paroxysmal events that are associated with hyperexcitability.

Together, our initial studies showed that BBB breakdown and astrocytic TGFβ signaling occur at early stages of mouse brain aging, concurrent with hyperexcitability and neural network dysfunction. Next, we performed comparative human studies to investigate whether similar BBB dysfunction and TGFβ signaling is also present in aging human subjects.

### The aging human brain shows progressive BBB dysfunction associated with astrocytic TGFβ signaling

In our rodent studies, we directly measured the progression of albumin extravasation and TGFβ signaling across the lifespan. BBB dysfunction has also been reported in aging humans including mild cognitive impairment (MCI) and Alzheimer’s disease and related dementia (ADRD) patients, using indirect measures such as CSF sampling or imaging, which can be technically challenging and may produce conflicting results *(15)*. Thus, we sought to further quantify age-related BBB dysfunction in human subjects, expanding on results reported elsewhere *(8, 10, 12, 13)*. We used dynamic contrast-enhanced MRI scanning (DCE-MRI) to quantify BBB permeability in 113 healthy human subjects with an age range of 21 to 83 years old (Fig. 2A; Table S1). We first established normal levels of brain permeability based on permeability values in healthy, young patients (age 21-40), setting an upper value for “normal permeability” at the 95^th^ percentile (i.e. 95% of brain voxels in the averaged healthy young brain were below this value). Based on this threshold, we constructed permeability maps for each subject and quantified the percentage of “leaky” voxels in the whole brain (Fig. 2A). This analysis revealed a significant linear increase in BBB permeability in aging (Fig. S2A). To estimate the overall prevalence of BBB dysfunction, we further categorically classified individuals as either BBB-intact (BBB-I) or BBB-disrupted (BBB-D; permeability in more than 5% of brain volume). By age 60, nearly half of the population was affected by BBB-D (Fig. S2B).

**Fig. 2.**
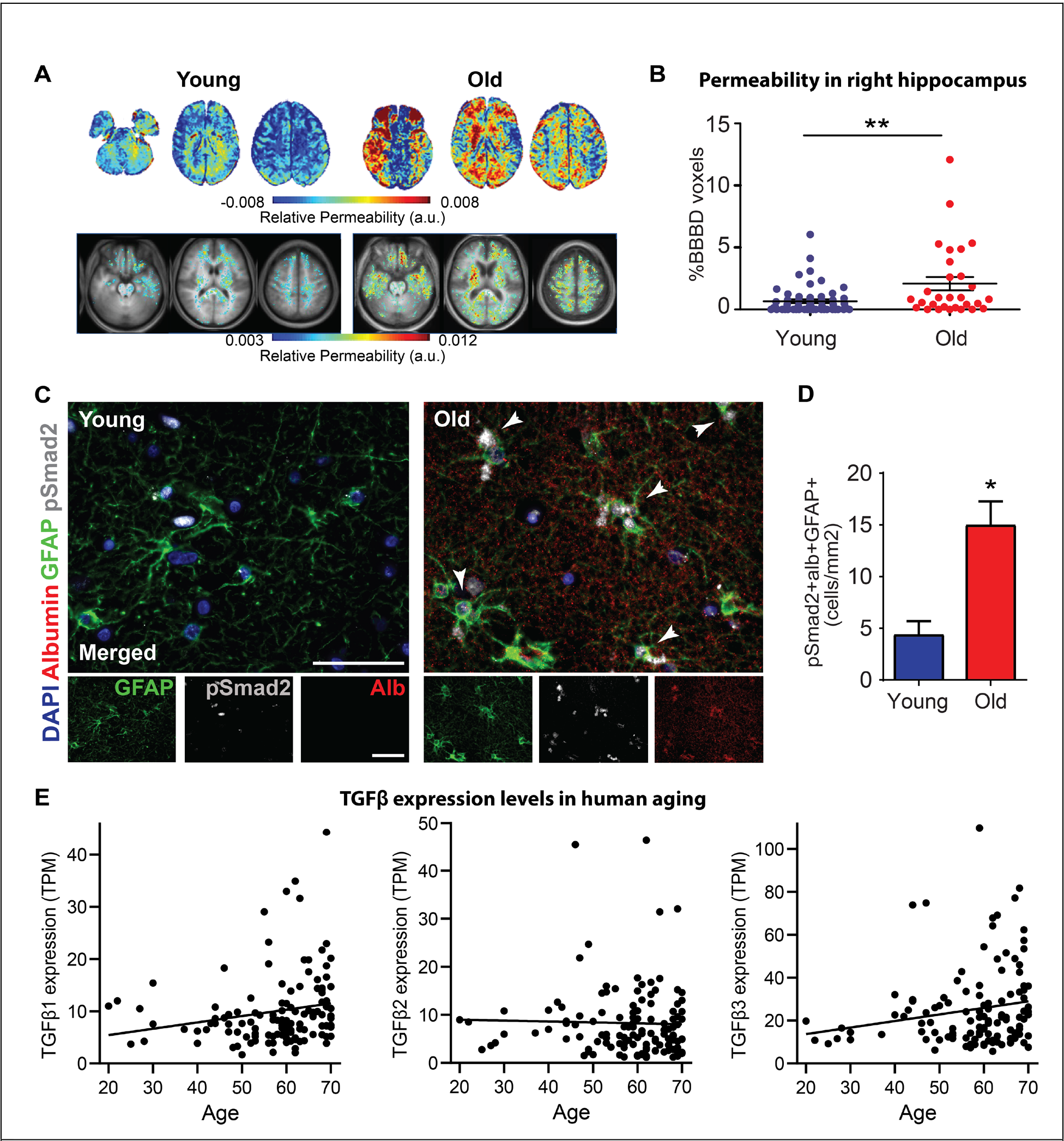
Aging patients show decline in the integrity of the BBB and increased TGFβ signaling. (A) (Top Panel): Representative images from DCE-MRI scans of a young (30 years old) and old (70 years old) subject using an MRI-sensitive contrast agent, Gd-DTPA, which does not cross the intact BBB. Intensity of signal reflects relative BBB permeability (in arbitrary units, a.u.). (Bottom Panel): Average permeability maps from all young (ages 20-40) and old (60–80) subjects. (B) BBB permeability was significantly elevated in the right hippocampus of old subjects compared to young. (C) Representative images of immunofluorescent staining of post-mortem human hippocampal tissue from young (31.3 ± 5 years) and old (70.6 ± 5.6 years) individuals, showing albumin in the aging hippocampus, localized in astrocytes (GFAP) and co-localized with pSmad2, indicated by arrows. Nuclei are visualized by DAPI stain. Scale bar = 50 μm. (D) The aging human hippocampus has significantly elevated numbers of astrocytes that colabel with albumin and pSmad2 (t-test, p=0.0324, n = 3 young, 10 old). (E) Scatterplots and best-fit linear regressions of TGFβ expression levels (transcripts per million; TPM) from hippocampus of human subjects aged 20-70 years old (n=123 subjects), accessed from the GTEx project expression database. Aging is significantly correlated with increasing hippocampal expression of TGFβ1 (Pearson correlation, r = 0.195, p = 0.031) and TGFβ3 (r = 0.184, = 0.042) isoforms, but not TGFβ2 (r = −0.028).

BBB breakdown, specifically in the hippocampus, has been associated with the incidence of mild cognitive impairment *(13)*. Furthermore, hippocampal atrophy and in particular asymmetrical atrophy are early biomarkers that predict the transition from cognitively normal to MCI to ADRD *(50–55)*. Thus, we used anatomical localization to specifically assess BBB permeability in the left and right hippocampus, and found that aging individuals showed asymmetric BBB breakdown localized to the right hippocampus (Fig. 2B). Given that our study was comprised of cognitively “normal” aging individuals, excluding subjects that met the diagnostic criteria of mild cognitive impairment, this suggests that BBB breakdown may be a very early aging event that precedes cognitive decline and neuropathology.

We next sought to complement the indirect MRI approach by directly assessing BBB dysfunction, and its association with astrocytic TGFβ signaling in aging human brains. We examined post-mortem tissue from young (31.3 ± 5 years old) and aging (70.6 ± 5.6 years) human subjects with no history of brain disorder (Table S2), and directly measured co-localization of albumin extravasation and TGFβ signaling, in astrocytes as performed in our rodent studies. We found high levels of the serum protein albumin in the old hippocampus, which was absent in the young (Fig. 2C). Albumin was detected in astrocytes (identified by the astrocytic marker GFAP), and colocalized with phosphorylated Smad2 (pSmad2), the primary signaling protein of the canonical TGFβ signaling cascade. The number of albumin-pSmad2 co-labeled astrocytes was significantly increased in old vs. young individuals (Fig. 2D).

This immunostaining indicated that BBB dysfunction and albumin induces TGFβ signaling in the aging human hippocampus, as observed in our aging rodent studies. In turn, this would be predicted to cause positive feedback and elevated levels of TGFβ. To assess this, we obtained human brain transcriptome data from the publicly available Genotype-Tissue Expression (GTEx) project *(56)*, with data from 123 human subjects ranging in age from 20 to 70 years old, and investigated brain expression levels of the major TGFβ isoforms (TGFβ1, 2, and 3) in the human hippocampus. These data showed that hippocampal expression of TGFβ1 and TGFβ3 significantly increase with age in human subjects (Fig. 2E).

These studies showed that BBB dysfunction and TGFβ signaling are early biomarkes of aging in both rodents and humans, and in rodents are associated with symptomatic hyperexcitability. Next, we used gain-of-function and loss-of-function interventions to determine whether albumin-induced TGFβ signaling plays a causal role in age-related symptoms of neural dysfunction and cognitive impairment

### Infusion of albumin into young brains causes hyperexcitability and paroxysmal slow wave events

To test if TGFβ signaling, as induced by BBB dysfunction, is sufficient to cause symptomatic pathology associated with aging, we infused albumin (iAlb) or control artificial cerebrospinal fluid (aCSF) into the brain ventricles of healthy, young adult rats and mice via osmotic mini-pump (Fig. S3A). Following infusion, the exogenous albumin diffused readily into the ipsilateral hippocampus, and was taken up by astrocytes within 48 hours of infusion (Fig. S3A-B). We then assessed outcomes in young rodents (Fig. 3A), to determine if iAlb is sufficient to cause aged-like symptoms of network and cognitive dysfunction. We conducted the hyperexcitability PTZ seizure assay in young mice 48 hours after iAlb. The iAlb mice showed significantly increased seizure severity and mortality induced by PTZ compared to aCSF controls (Fig. 3B-C; Fig. S3C), fully reprising the hyperexcitable seizure vulnerability observed in naturally aged mice. Furthermore, we directly investigated network dysfunction via recordings of ECoG activity in young iAlb rats, and found that iAlb caused a symptomatic slowing of ECoG activity and a significantly increased pSWEs (Fig. S3D-G; Fig. 3D) similar to the aberrant activity observed in aged rodents. This elevated level of pSWEs was observed only in recordings from the ipsilateral hemisphere receiving iAlb infusion, but not in the contralateral hemisphere, indicating specificity of the aberrant neural activity to the tissue affected by iAlb.

**Fig. 3.**
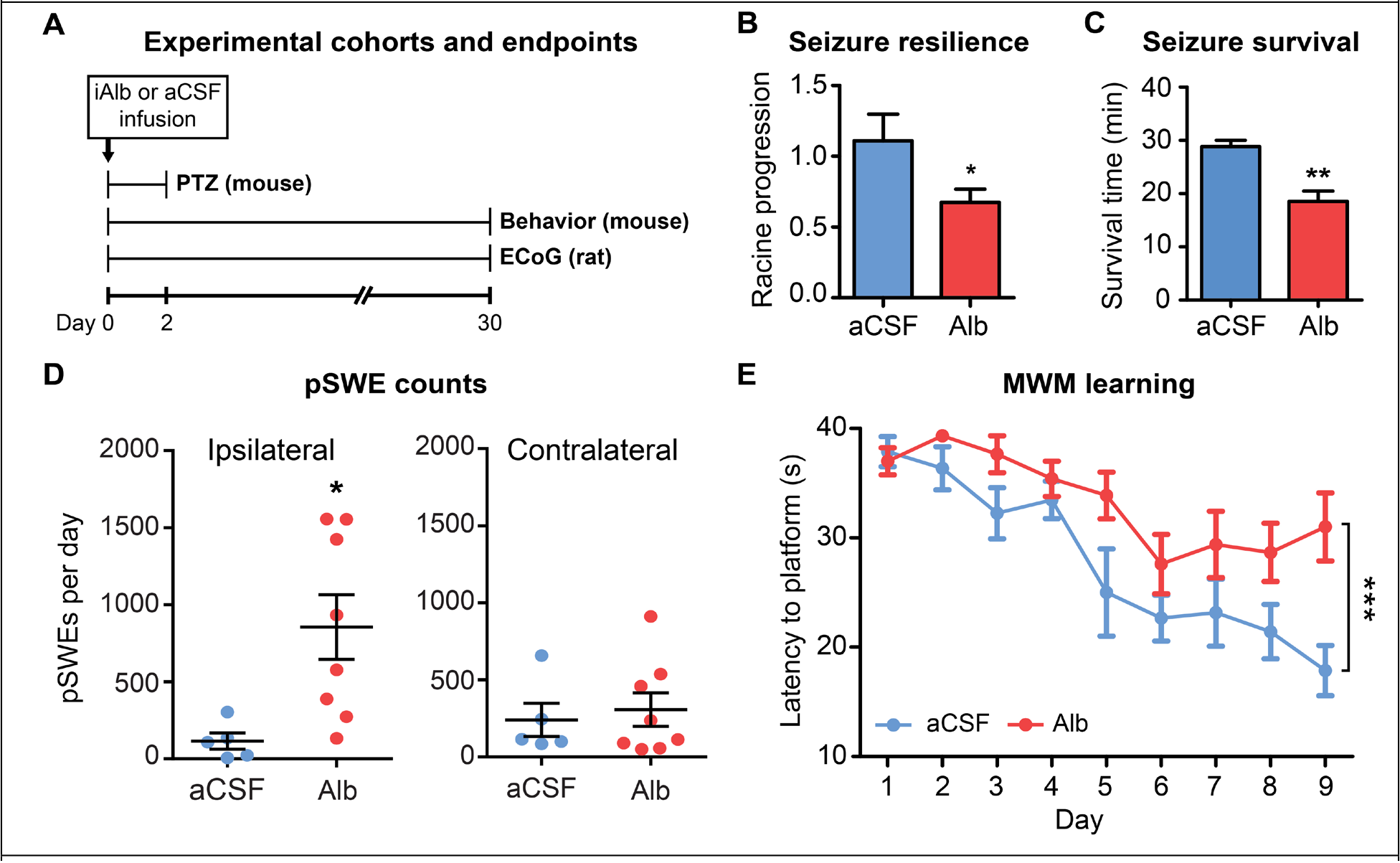
Induction of TGFβ signaling in young rodents causes aberrant network activity, vulnerability to induced seizures, and cognitive impairment. (A) Timeline of outcome measures (endpoints) taken in cohorts of young adult rats and mice after icv albumin infusion (iAlb). (B) Young mice received iAlb or aCSF control infusion for 48 hours, followed by PTZ seizure induction. Albumin infusion caused increased severity in induced seizures with regression slopes showing faster seizure progression (t-test with Welch’s correction, p = 0.0497, n = 4), and (C) faster latency to mortality (t-test, p=0.004), compared to aCSF controls. (D) ECoG activity was recorded from rats following one month of iAlb infusion. iAlb rats had significantly elevated numbers of pSWEs compared to aCSF-infused controls in the ipsilateral hemisphere receiving infusion, but not the contralateral hemisphere. (E) Young mice were given iAlb surgery and then tested in the MWM task one month later. Compared to aCSF controls, iAlb mice had significantly poorer learning performance (repeated measure ANOVA, n = 9 aCSF, 10 iAlb, main effect of learning over time p < 0.0001, main effect of group p = 0.0093, with Bonferroni posttest on day 9 showing significant differences between aCSF and iAlb, p < 0.001). For all tests, *p<0.05, **p<0.01, ***p<0.005, ****p<0.001.

### Infusion of albumin into young brains causes impaired memory performance

We next investigated whether iAlb is sufficient to cause cognitive impairment in young mice. We implanted mice with iAlb (or aCSF control) osmotic pumps for one week. One month later, we tested mice in the Morris water maze (MWM) spatial memory task. Mice that received iAlb infusion were significantly impaired in memory performance over 9 days of MWM training, compared to control mice with aCSF implant (Fig. 3E). Together, these data show that iAlb is sufficient to induce TGFβ signaling and confer a dramatic “old age” phenotype in young rodents, including aberrant neural ECoG activity, hyper-excitable seizure vulnerability, and cognitive impairment.

### Genetic knockdown of astrocytic TGFβ signaling reverses pathological outcomes in aging mice

To test the causal role of astrocytic TGFβ signaling in age-related impairments, we generated a transgenic mouse line (aTGFβR2/KD) expressing inducible Cre recombinase under the astrocyte-specific GLAST promoter. This enables conditional knockout of the floxed (fl) TGFβR with temporal precision, specifically in astrocytes (Fig. 4A), allowing us to interrogate the role of astrocytic TGFβ signaling in mediating pathological outcomes. Treatment with tamoxifen (tam) induced efficient recombination in approximately 40% of hippocampal astrocytes but not in neurons (Fig. S4A-C), and significantly reduced levels of TGFβR (Fig. S4D), thus effectively causing knockdown (KD) to inhibit but do not fully abolish TGFβ signaling. We aged cohorts of aTGFβR2/KD mice to early (12-16 months) and late (17-24 months) stages of aging, and then induced TGFβR2 KD in astrocytes to test their role in age-related hyperexcitability and cognitive dysfunction (Fig. 4B). Following induction, aged KD mice showed a significant decrease in hippocampal pSmad2 protein and reduction in expression of TGFβR2 (Fig. S4E-F).

**Fig. 4.**
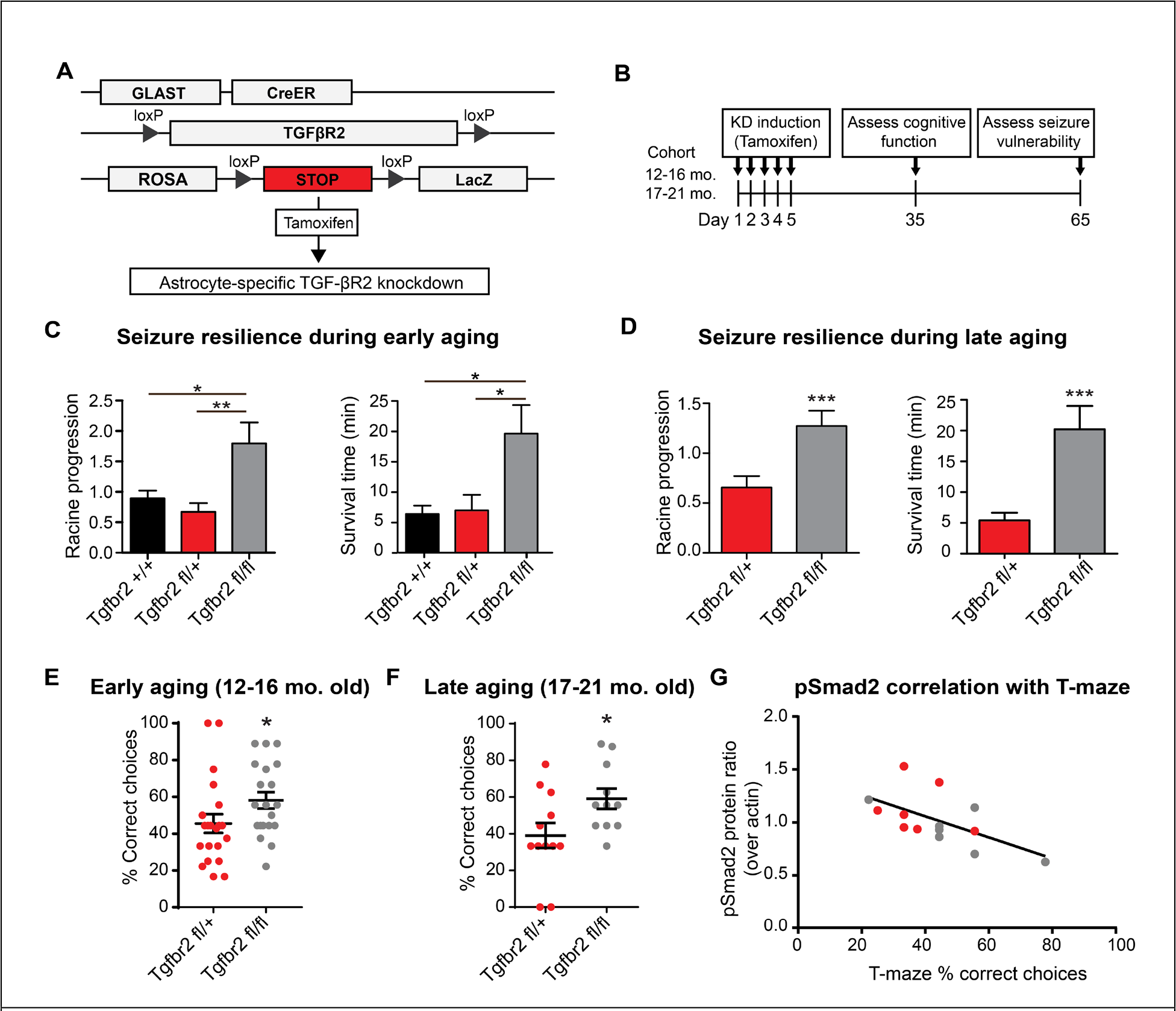
Knockdown of astrocytic TGFβ reverses neurological outcomes in aging mice. (A) Schematic of the transgenic aTGFβR KD system. The astrocytic GLAST promoter drives expression of Cre recombinase in astrocytes. Following induction with tamoxifen injection, activated Cre excises the TGFβR gene at inserted floxed (fl) loxP sites. LacZ reporter expression provides a readout of Cre activity. (B) Experimental timeline. KD was induced in early (12-16 mo.) and late (17-24 mo.) aged mice, and T-maze testing was performed 35 days later. At 65 days post-induction, mice were tested for vulnerability to PTZ-induced seizures. (C) aTGFβR KD (Tgfbr2^fl/fl^) reversed seizure vulnerability in 12-16 mo old mice, significantly slowing progression through the Racine scale (1-way ANOVA, p = 0.0019, with Bonferroni posttest, n = 5 Tgfbr2^+/+^, 6 Tgfbr2^fl/+^ and Tgfbr2^fl/fl^) and latency to mortality (1-way ANOVA, p = 0.022, with Bonferroni posttest), compared to control heterozygous mice (Tgfbr2^fl/+^) and mice given oil control rather than tamoxifen induction (Tgfbr^+/+^). (D) 17-24 month old mice with aTGFβ KD were similarly protected against PTZ seizure vulnerability with less severe seizures (t-test with Welch’s correction, p = 0.002, n = 5 Tgfbr2^fl/+^ and 9 Tgfbr2^fl/fl^) and delayed mortality (t-test with Welch’s correction, p = 0.004). (E-F) In T-maze, aTGFβR KD mice showed significantly better working memory performance compared to heterozygous controls at in both early aging (12-16 months old, Mann-Whitney test, p = 0.0273, n = 21 Tgfbr2^fl/+^, 20 Tgfbr2^fl/fl^) and late aging assessments (t-test, p = 0.035 n = 12 Tgfbr2^fl/+^, 11 Tgfbr2^fl/fl^). (G) An additional cohort of 12-16 month old mice was tested in T-maze, and hippocampi were dissected for Western blot analysis of TGFβ signaling (pSmad2) to assess individual outcomes. T-maze performance was negatively correlated with levels of pSmad2 signaling across individuals of both genotypes (Tgfbr2^fl/+^ and Tgfbr2^fl/fl^) (Pearson’s correlation, r = −0.598, p = 0.024, n = 14). For all tests, *p<0.05, **p<0.01, ***p<0.005, ****p<0.001.

In early aging (12-16 months), mice with homozygous induced KD of the aTGFβR2 (fl/fl) were protected against symptomatic hyperexcitability, showing a low level of vulnerability to PTZ-induces seizures and seizure mortality (Fig. 4C; Fig. S4G) that was similar to young mice. In contrast, 12-16 month old control mice that were heterozygous for floxed TGFβR2 (fl/+) (and hence retained intact astrocytic TGFβ signaling), or that were homozygous for floxed TGFβR2 but injected with vehicle instead of tamoxifen (hence no KD induction; aTGFβR2 +/+) showed significantly elevated levels of seizure vulnerability and mortality (Fig. 4C). The KD intervention was also effective at late aging timepoints: 17-21 month aged aTGFβR2 heterozygote (fl/+) controls showed typical age-related vulnerability to induced seizures, whereas aTGFβR2 KD (fl/fl) displayed low vulnerability and mortality to PTZ challenge (Fig. 4D; Fig. S4H). These results show that genetic KD of astrocytic TGFβ signaling is sufficient to reverse symptoms of hyperexcitability in the PTZ assay, at both early and late stages of aging.

### Genetic knockdown of astrocytic TGFβ signaling reverses cognitive impairments in aging mice

To assess the role of astrocytic TGFβ signaling in age-related cognitive decline, we tested aged transgenic mice for spontaneous alternation in the T-maze, a hippocampal working memory task *(57)* that is impaired in aging rodents *(58)*. The task is optimal for assessing aging rodents because it can be performed rapidly without extensive training, is sensitive to mild impairments in hippocampal function *(59, 60)*, and yet it is relatively unaffected by motor and vision impairments that may confound aging mice in traditional tasks such as Morris Water maze *(57)*. At both early and late aging time points, aTGFβR2 KD mice made significantly more correct choices in the T-maze task, indicating improved working memory compared to heterozygous controls (Fig. 4E-F).

While aTGFβR2 KD was effective at improving cognitive function in both early and late stages, we also found greater heterogeneity in cognitive scores in the early (12-16 month) aging group – as would be expected for an early stage of aging in which some mice may have transitioned into mild cognitive impairment while other remain cognitively healthy. To investigate this individual variability, we performed T-maze in an additional cohort of early aging (12-16 month old) aTGFβR2 KD and heterozygous controls, and collected dissected hippocampi to quantify the relationship individual cognitive scores and pSmad2, the molecular marker of TGFβ signaling. Across both heterozygous (fl/+) and homozygous (fl/fl) genotypes, pSmad2 levels were negatively correlated with T-maze performance (Fig. 4G), providing further evidence for the role of TGFβ signaling in cognitive impairment.

Together, these results show that targeted inhibition of the TGFβ signaling pathway, via induced KD in astrocytes, is sufficient to reverse the outcomes of seizure vulnerability and cognitive impairment in a hippocampal spatial working memory task in old mice, and that cognitive outcomes in a heterogeneous “mildly impaired” early aging cohort are correlated with the individual levels of TGFβ signaling.

### A novel small molecule TGFβR1 kinase inhibitor blocks iAlb-induced effects in the young brain

Our findings in mice and human brains support an evolutionarily conserved role of astrocytic TGFβ signaling in the pathogenesis of age-related neurological vulnerability, further indicating a therapeutic potential in targeting TGFβR. Thus, we next tested the efficacy of a novel small molecule TGFβR1 kinase inhibitor, IPW *(61)*. IPW has a promising clinical profile – including the ability to cross the BBB and good stability following oral dosing (Fig. S5A), making it suitable for once-per-day dosing to achieve inhibition of TGFβR signaling (Rabender et al., 2016). We first tested IPW in young mice with TGFβ signaling induced by iAlb, treating them with daily i.p. injections (20 mg/kg) of IPW for two days after pump implant. IPW treatment significantly reduced pSmad2 levels measured in the dissected hippocampus of treated mice, compared to mice treated with vehicle control (Fig. 5A-B). When examining the vulnerability to PTZ, iAlb mice treated with vehicle showed high seizure vulnerability, replicating our previous results, whereas IPW treatment effectively reversed this vulnerability, reducing both seizure severity and mortality to the level seen in the control aCSF infused mice (Fig. 5C-D; Fig. S5B). These data supported the likelihood for IPW efficacy in naturally aged mice, since it showed not only excellent target engagement, but also efficacy in reducing symptomatic hyperexcitability induced by iAlb.

**Fig. 5.**
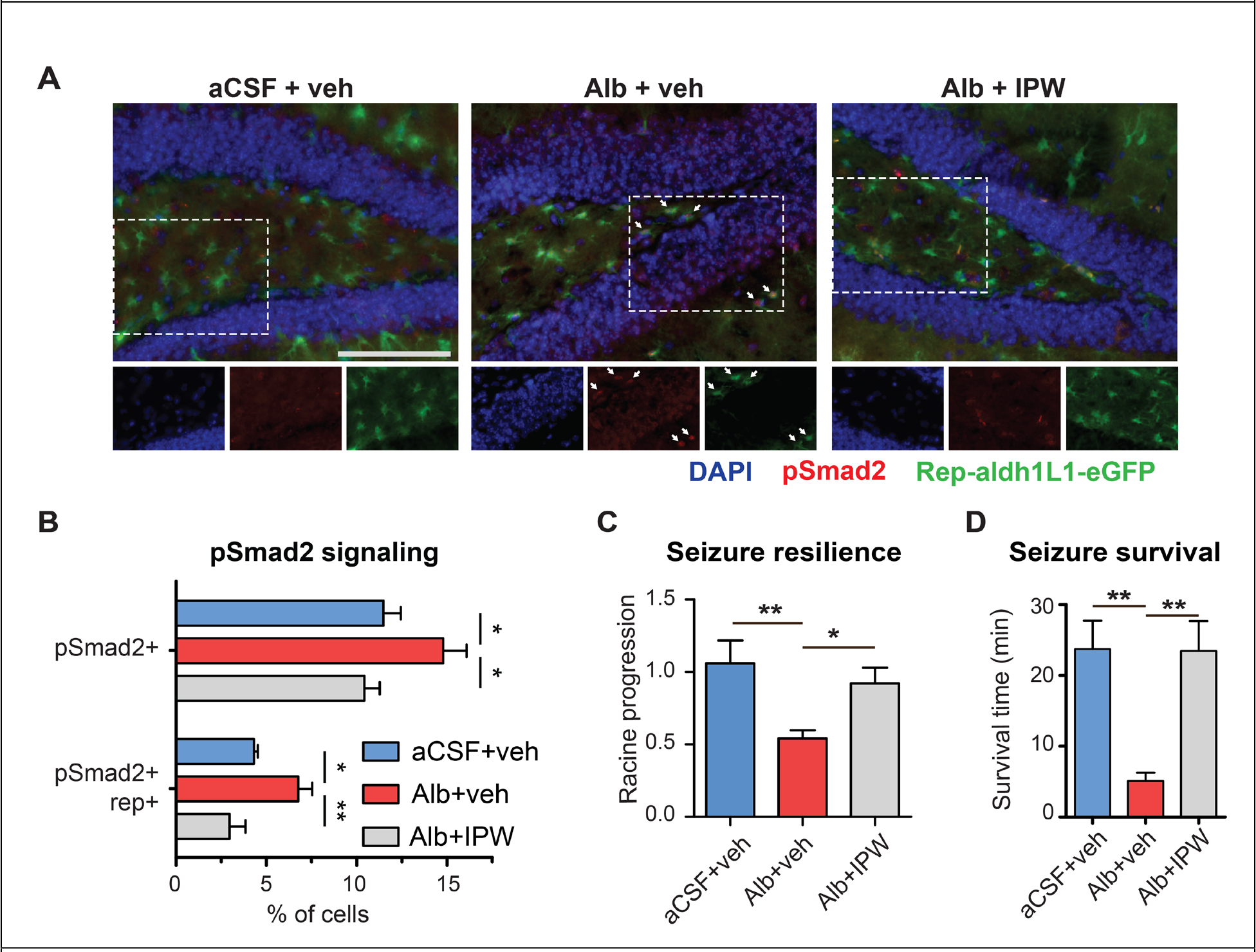
IPW reduces TGFβ signaling and seizure vulnerability in young mice infused with albumin. (A) Immunofluorescent images from iAlb mice after treatment with IPW or vehicle injections during 7 days of icv albumin infusion (Scale bar = 100 μm; dotted box indicates region of inset image; arrows indicate examples of astrocytes colabeled with pSmad2). (B) iAlb mice showed elevated levels pSmad2 signaling localized in astrocytes (compared to aCSF controls) and IPW reduced pSmad2 (1-way ANOVA with Neuman-Keuls Multiple Comparison Test: % pSmad2, p = 0.027; % pSmad2+ rep+, p = 0.0048; n = 6). (C) IPW treatment reduced seizure vulnerability conferred by iAlb, restoring seizure profile to aCSF control levels (1-way ANOVA, main effect of treatment p = 0.004, with Bonferroni’s posttest, n = 6 (aCSF+veh and Alb+veh), 7 (Alb+IPW)). (D) IPW treatment also reversed the vulnerability to seizure mortality conferred by iAlb (1-way ANOVA with Tukey’s posttest, p=0.003, n = 6 (aCSF+veh and Alb+veh), 7 (Alb+IPW). For all tests, *p<0.05, **p<0.01, ***p<0.005, ****p<0.001.

### Drug inhibition of TGFβ signaling reverses molecular and functional brain aging in mice

Based on these validation studies, we then tested IPW as an intervention against TGFβ signaling in mice aged to 2 years old, near the end of the mouse lifespan. Immunofluorescent analysis of hippocampal sections from aged mice revealed that 5 days of treatment with IPW (20 mg/kg i.p.) reduced the number of astrocytes co-labeled with pSmad2 (Fig. 6A). Similarly, hippocampal Western blot showed that 5 days of IPW treatment reduced the high levels of pSmad2 in aged mice, thus restoring a “healthy” level of TGFβ signaling similar to that of young mice (Fig. 6B). Furthermore, IPW treatment reduced the downstream output TGFβ1 (Fig. 6C).

**Fig. 6.**
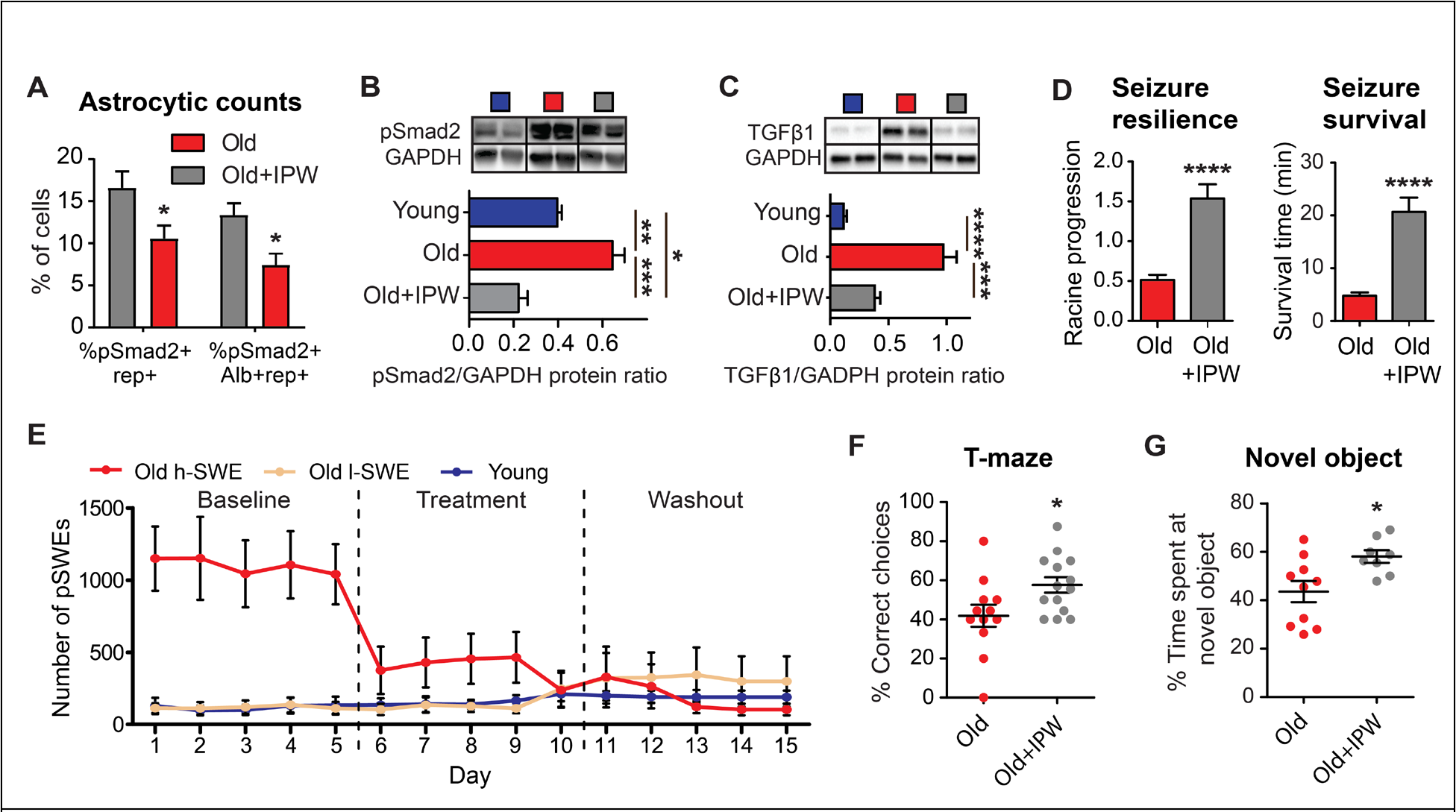
IPW reverses TGFβ signaling, aberrant neural activity, seizure vulnerability, and cognitive impairment in aged mice. (A) Aged Rep-Aldh1L1 mice were treated 7 days of IPW (20 mg/kg, i.p.) or vehicle, followed by immunofluorescent staining for pSmad2 and albumin (Alb). IPW significantly reduced fraction of astrocytes (rep+) that co-labeled with pSmad2 (t-test, p=0.037), and the fraction of cells triple labeled with pSmad2, Alb, and rep (t-test, p=0.014) (n = 5 old, 6 old+IPW). (B) Densitometry of Western blot shows that IPW restored levels of elevated pSmad2 protein in old mice, to the level of young mice (ANOVA, main effect p=0.0001, with Bonferroni posttest, n = 4). IPW treatment also reduced levels of (C) TGFβ1 in aged mice (ANOVA, main effect p<0.0001, with Bonferroni posttest, n = 4). (D) 7 days of IPW treatment reversed seizure vulnerability in aged mice, reducing progression through the Racine scale (t-test, p < 0.0001, n = 9) and reducing the onset of seizure induced mortality (t-test, p <0.0001). (E) Continuous ECoG recording over 15 days was used to investigate effects of IPW on aberrant network activity. During 5 days of baseline recording, clustering analysis was used to classify symptomatic old mice as affected by high numbers of pSWEs (h-SWE, n = 6), or asymptomatic old mice with low numbers of pSWEs (l-SWE, n = 6) that were similar to the level of young mice (n = 5). In the subsequent 5 days of recording, IPW treatment restored a youthful ECoG profile in the affected mice, reducing pSWEs (Dunn’s multiple comparisons test p = 0.0098) to the level of young mice. After IPW dosing was halted, the old mice retained an asymptomatic ECoG profile throughout the “washout” period (Dunn’s multiple comparisons test baseline vs washout p = 0.0002). 7 days of IPW treatment improved cognitive scores of old mice in the (F) T-maze (t-test, p = 0.028, n = 12 vehicle, 14 IPW) and (G) novel object memory tasks (t-test, p = 0.017, n = 10 vehicle, 8 IPW). For all tests, *p<0.05, **p<0.01, ***p<0.005, ****p<0.001.

To test if IPW inhibition of TGFβ signaling could reverse the symptoms of neural hyperexcitability in aged mice, we treated 24-month-old mice with IPW for 7 days, and then performed the PTZ assay. As in aTGFβR KD genetic intervention, mice treated with IPW showed lower seizure severity and mortality compared to aged control mice treated with vehicle (Fig. 6D; Fig. S6A). To further investigate the efficacy of IPW on aberrant neural activity, we conducted a longitudinal experimental design in which young and old mice from the ECoG cohort were continuously recorded for 5 days of baseline, followed by 5 days of IPW treatment, and then 5 days of “washout” (no further dosing). This design was intended to assess not only the acute efficacy of IPW for treating symptoms of aberrant ECoG activity, but also whether any treatment effects persist after dosing is halted.

During the baseline recording period, we observed that approximately half of the aged mice showed a phenotype of high pSWEs, whereas others showed no aberrant ECoG activity. These subgroups were confirmed by unbiased clustering analysis, which defined a best fit for the aged mouse population as two distinct clusters of high pSWE (h-SWE) and low pSWE (l-SWE) mice (Fig. S6B-D). In the longitudinal experimental design, we thus analyzed these subgroups separately, in order to assess the efficacy of IPW treatment on the h-SWE phenotype. In h-SWE mice, treatment with IPW dramatically reduced the number of pSWEs, restoring a profile of ECoG activity similar to that of young mice (Fig. 6E). In contrast, IPW treatment had no effect on l-SWE mice, or on young mice, which did not show any aberrant high pSWE activity. Furthermore, treatment of aged mice with vehicle control showed no efficacy on reducing pSWEs (Fig. S6E-F). Beyond the treatment phase, the efficacy of IPW on reducing h-SWEs also persisted through the end of the washout period, indicating that inhibition of TGFβ signaling may mediate a long-lasting change in the underlying hyperexcitability of neural circuits.

Given the effects of IPW on inhibiting TGFβ signaling and reversing associated outcomes of hyperexcitability and ECoG neural network dysfunction, we next assessed functional cognitive outcomes. Aged mice were treated for 7 days with IPW or vehicle control, and then assessed in two cognitive behavioral tasks performed over consecutive days: spontaneous alternation in T maze, and the novel object task, which is also sensitive to age-related memory decline *(62, 63)*. After 7 days of IPW treatment, aged mice showed significant improvement in both cognitive tasks, relative to vehicle-treated controls (Fig. 6F-G), demonstrating that IPW is effective in improving cognitive impairment in aged mice. Together, these studies show that IPW inhibition of chronic TGFβ signaling in aged mice can rapidly restore a “youthful” profile of network activity and cognitive capacity.

## Discussion

Aging is often accompanied by cognitive decline, even in the absence of dementia or measurable neurodegeneration *(64, 65)*. Unlike transgenic models for artificially inducing age-like disease, our investigations focused on naturally aging mice, allowing us to observe the relative sequence of biological changes associated with brain aging. We found that BBB dysfunction and consequent albumin extravasation appears as early as middle age, placing it among the earliest known hallmarks of the aging brain. Consistent with our findings, relatively subtle changes in neural and synaptic function have been widely observed in humans and other mammals as one of the first signs of neurological aging *(64–67)*, and these changes in neurotransmission are associated with hippocampal hyperexcitability that is thought to be one of the earliest events in the progression of mild cognitive impairment *(43, 44, 68)*. However, the regulatory pathways that may trigger or control these changes are unknown. We found that microvascular BBB dysfunction allows for the extravasation of serum albumin into the brain and hyperactivation of TGFβ signaling, similar to the activation of TGFβ signaling that has been shown in head injury models *(22)*. Activation of this signaling cascade in turn causes symptoms associated with aberrant neural function (Fig. 7).

**Fig. 7.**
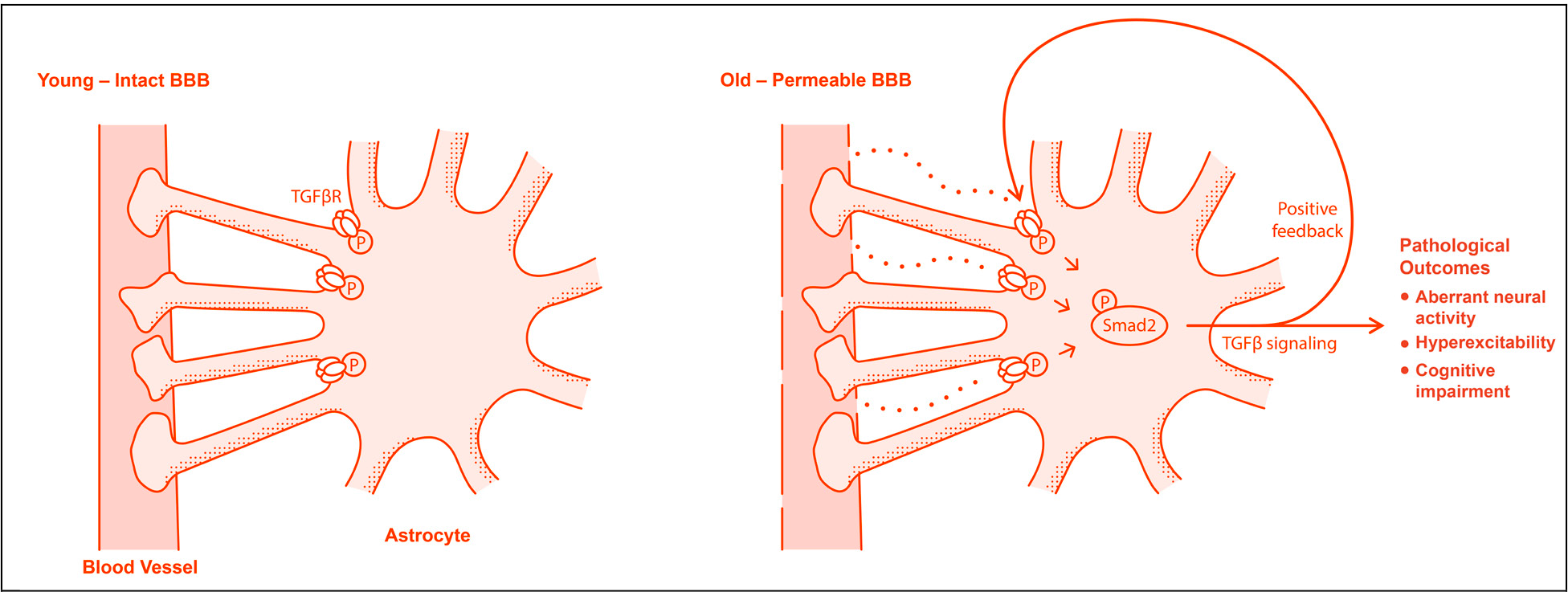
Schematic summarizing the mechanistic model underlying BBB-related neural pathology. In old age, serum proteins including albumin cross the dysfunctional BBB and activate astrocytic TGFβ receptor (TGFβR) signaling. Outputs of the astrocytic TGFβ signaling cascade include positive feedback and induction of symptomatic pathology including aberrant neural activity, hyperexcitability, and cognitive impairment.

We used telemetric ECoG to directly record abnormal neural network activity during aging. We found slowing of EEG/ECoG activity, consistent with other reports in the context of aging *(46–48)*. We further showed that this slowing of activity in mice is characterized as discrete, paroxysmal transient events (pSWEs), occuring frequently and spontaneously against a backdrop of “normal” ECoG activity. These pSWEs, which have characteristics that are similar to but distinct from epileptic seizures, may constitute “silent” or “subclinical” epileptiform activity, as has been reported in human dementia patients *(69–71)*. These data support the suggestion, proposed elsewhere *(72–74)*, that there may be common mechanistic links between age-related dementia and epilepsy, which has a remarkably high incidence in the elderly *(75–77)*. We emphasize that elevated TGFβ signaling, triggered by BBB dysfunction, provides a simple, parsimonious model for how this dysfunction may arise in aging. Indeed, we found that TGFβ inhibition reversed aberrant ECoG activity, increased seizure threshold, and improved cognitive outcomes in aged mice, suggesting efficacy in the myriad symptoms that would be expected to arise from a dysfunctional neural network.

Could inhibition of TGFβ signaling, as a strategy to counteract the detrimental consequences of age-related BBB dysfunction, hold therapeutic potential? One of the major challenges (and causes of failure) in treating progressive neurological diseases is that patients decline over time, and thus may accumulate irreversible damage by the time of diagnosis. Considering that BBB dysfunction begins relatively early in aging, this might call for a scenario in which chronic preventive treatment with TGFβ inhibitors is requried to avoid future damage. However, we found that one week of acute treatment reversed the pathological outcomes in aged mice, including elevated TGFβ signaling, aberrant ECoG activity, seizure vulnerability, and cognitive dysfunction. Our findings suggest that the aging brain may retain considerable cognitive capacity, which may be chronically suppressed (but not irreversibly lost) by BBB leakiness and its inflammatory fallout.

By uncovering a foundational mechanism linking BBB decline to neural dysfunction, our study raises several critical questions: what causes BBB decline itself? And is BBB decline causal to, concurrent with, or an outcome of (or independent from) other well-known mechanisms of aging such as inflammation, reactive oxidation stress and metabolic failures, proteasome senescence, DNA damage, etc. *(78–83)*? Our work, while novel in focusing on microvascular integrity and interactions within the neurovascular unit, can be placed in context of some of these mechanisms. For example, activation of astrocytes and gliosis has been shown to be a key step in many different aging diseases, with astrocytes playing potent roles in controlling neuroinflammation and neural functions including synaptic plasticity, senescence, and neurodegeneration *(84–88)*; we show that BBB dysfunction may be an early step causing or contributing to activation of astrocytes and the ensuing inflammatory response. Similarly, several previous studies have reported increased brain TGFβ signaling in aging *(89–94)*, and suggested that it could be a primary regulatory factor inducing aged neural phenotypes – although it was unknown what may trigger this increase in TGFβ signaling in aged individuals. We show that extravasation of albumin through the leaky BBB and astrocytic uptake may be one of the earliest steps that induces this age-related TGFβ cascade. Ultimately, our mechanistic findings provide a guiding framework for translation into the human clinical context, including large-scale epidemiological studies that are needed to establish the relationship between BBB status, other known biomarkers, and disease outcomes. In particular, this research offers new hope in two key unmet areas: early detection (via MRI imaging of BBB status), and a new avenue for disease-modifying treatment that is mechanistically distinct from other canonical dementia targets, many of which have failed in clinical trials.

## Materials and Methods

### Study design and statistical analysis

The aim of this study was to use experimental gain-of-function (iAlb) and loss-of-function (TGFβR KD and IPW) interventions to investigate the causal role of TGFβ signaling in age-relate BBB pathology and the efficacy of therapeutic intervention. Outcomes were assessed using molecular (immunofluorescent staining and WB), electrophysiological (ECoG), and behavioral (PTZ and cognitive tasks) measures of symtomatic pathology in rodents. Translatability of these findings was further supported by complementary measures of BBB permeability and TGFβ signaling in human subjects (via CTE-MRI imaging and post-mortem histology). The experimental designs and methodology of outcome measures were chosen prior to initiating each experiment, except in the case of the ECoG experiments, in which post-hoc analysis of ECoG signal led to the discovery of pSWEs as a novel biomarker of aberrant network activity. For all experiments, subjects were randomly assigned to experimental groups, and data were collected under blinded experimental conditions with the exception of the immunofluorescent albumin counts in Fig. 1A, in which differences between age groups were visually apparent in the tissue even under naïve conditions, such that complete blinding was not possible. The lead and corresponding authors were responsible for experiment design, conducting statistical analysis, and unblinding the final results. All graphs are plotted showing mean and SE. Two sample comparisons were performed by student’s t-test or Mann-Whitney test, and multiple group comparisons were conducted by ANOVA or Kruskal-Wallis test, followed by post-hoc testing to compare individual groups when a main effect was detected. Multiple correction comparisons were used as described in figure legends. Seizure progression in the PTZ experiments was analyzed by two-way ANOVA, and linear regression was used to calculate regression slopes. Differences in regression slope were also compared by ANOVA. In mouse ECoG experiments, different subgroups (h-SWE and l-SWE) were observed after data collection. These subgroups were formally classified using an unbiased Gaussian mixed model, and then inferential statistics were performed on the subsequent groups. For all inferential statistics, two-tailed tests were used and significance thresholds were set at p<0.05. For rodent behavior experiments, animals were removed from the study cohort if they were unable to complete the requisite task, according to the following pre-defined criteria: for MWM, mouse shows inability to swim (unable to maintain swim speed or buoyancy); for T-maze, mouse does not leave stem to complete arm choice in greater than three trials. For TGFβ1 WB analysis following IPW treatment (Fig. 6C), two samples were excluded due to gel loading error. Exclusion of these data did not alter statistical significance (results of ANOVA analysis were significant with or without excluded values).

### BBB imaging

The human imaging protocol was approved by the Soroka University Medical Center Helsinki institutional review board, and written informed consent was given by all participants. BBB status was assessed by DCE-MRI in n=105 subjects with an age range of 21 to 83 years old. MRI scans were performed using a 3T Philips Ingenia scanner, and included: T1-weighted anatomical scan (3D gradient echo, TE/TR = 3.7/8.2 ms, acquisition matrix 432×432, voxel size: 0.5×0.5×1 mm), T2-weighted imaging (TE/TR = 90/3000 ms, voxel size 0.45×0.45×4 mm). For the calculation of pre-contrast longitudinal relaxation times (T10), variable flip angle (VFA) method was used (3D T1w-FFE, TE/TR = 2/10 ms, acquisition matrix: 256×256, voxel size: 0.89×0.89×6 mm, flip angles: 10, 15, 20, 25 and 30°). Dynamic contrast-enhanced (DCE) sequence was then acquired (Axial, 3D T1w-FFE, TE/TR = 2/4 ms, acquisition matrix: 192×187 (reconstructed to 256×256), voxel size: 0.9×0.9×6 mm, flip angle: 20°, ∆t = 10 Sec, temporal repetitions: 100, total scan length: 16.7 minutes). An intravenous bolus injection of gadoterate meglumine (Gd-DOTA, Dotarem, Guerbet, France) was administered using an automatic injector after the first five DCE repetitions. Data preprocessing included image registration and normalization to MNI coordinates (using SPM (http://www.fil.ion.ucl.ac.uk/spm)). BBB permeability was calculated for each brain voxel using in-house MATLAB script (Mathworks, USA), as described*(95–97)*. Briefly, a linear fit is applied to the later part of the concentration curve of each voxel; the slope is then divided by the slope at the superior sagittal sinus, to compensate for physiological (e.g., heart rate, blood flow) and technical (e.g., contrast agent injection rate) variability. For region specific permeability, disrupted region voxels were divided by total region voxels according to SPM registration.

### Immunostaining

For human staining, postmortem hippocampus was obtained from young (n=3, mean age = 31.3 ± 5 years) and old patients (n=10, mean age = 70.6 ± 5.6 years). All participants gave informed and written consent and all procedures were conducted in accordance with the Declaration of Helsinki and approved by the University of Bonn ethics committee. Resected hippocampi were fixed in 4% formalin and processed into liquid paraffin. All specimens were sliced at 4 μm with a microtome (Microm, Heidelberg, Germany), mounted on slides, dried, and deparaffined in descending alcohol concentration. For mouse samples, mice were anesthetized with Euthasol euthanasia solution and transcardially perfused with ice cold heparinized physiological saline (10 units heparin/mL physiological saline) followed by 4% paraformaldehyde (PFA, Fisher Scientific #AC416785000) in 0.1 M phosphate buffered saline (PBS). Brains were removed, post-fixed in 4% PFA for 24 hours at 4° C, and cryoprotected in 30% sucrose in PBS. Brains were then embedded in Tissue-Tek O.C.T. compound (Sakura, Torrance, CA), frozen, and sliced on a cryostat into 20 μm coronal sections, mounted on slides. Samples were immunostained under the following protocol. Slides were treated for antigen retrieval (for human, 5 min incubation at 100° C in sodium citrate buffer, pH 6.0); for mouse, 15 min incubation at 65 °C in Tris-EDTA buffer (10mM Tris Base, 1 mM EDTA solution, 0.05% Tween 20, pH 9.0), then incubated in blocking solution (5% Normal Donkey Serum in 0.1% Triton X-100/TBS) for 1 hour at room temperature. Samples were then stained with primary antibody at 4° C, followed by fluorescent-conjugated secondary antibody for 1 hour at room temperature, and then incubated with DAPI (900 nM; Sigma-Aldrich) to label nuclei. For human, primary antibodies were rabbit anti-phosphorylated Smad2 (Millipore AB3849-I, 1:500), chicken anti-Albumin (Abcam ab106582, 1:500), and mouse anti-GFAP (Millipore MAB3402, 1:500); for mouse, the same were used except anti-phosphorylated Smad2 (Millipore AB3849) and goat anti-GFAP (Abcam ab53554, 1:1000). Secondary antibodies were anti-rabbit Alexa Fluor 568, anti-chicken Alexa Fluor 647, anti-goat Alexa Fluor 488, anti-mouse Alexa Fluor 488 (1:500, Jackson ImmunoResearch), and anti-goat Alexa Fluor 647 (Abcam ab150131, 1:500). All antibodies dilutions were in blocking solution. For tissue from human patients and aged mice, slide-mounted brain sections for treated with TrueBlack Lipofuscin Autofluorescence Quencher (Biotium #23007) before coverslip mounting. Images were acquired at 20X or 40X objective magnification using a Zeiss Axio Observer Research microscope (Carl Zeiss AG) and Metamorph software (version 7.7.7.0), and analyzed using ImageJ software (NIH). For human samples, counts were performed in 10 of randomly selected sampling areas per subject. For mouse, imaging was performed in at least 3-4 hippocampi per mouse. Cell counts for each individual marker, and colabeling, were calculated manually by an observer, and normalized to the area or total number of DAPI-positive cells. For each subject, counts from each sampling area were averaged.

### Human transcriptome analysis

To investigate TGFβ gene expression patterns in brain hippocampus we evaluated expression data of 123 individuals from the Genotype-Tissue Expression (GTEx) project v7 release *(56)*. RNA sequencing was done by the GTEx consortium to measure the gene expression level from samples, reported as transcripts per million (TPM). We obtained data derived from hippocampal tissue samples for expression of the TGFβ isoforms TGFβ1 (ENSG00000105329.5), TGFβ2 (ENSG00000092969.7), and TGFβ3 (ENSG00000119699.3), and performed linear regression and Pearson correlations on the resulting datasets with respect to subject age.

### Evans Blue assay

4% Evans Blue (EB) solution in 0.9% sterile saline was administered by i.v. bolus injection through the tail vein at a dose of 2 mL/kg. Thirty minutes after tracer administration, brains were transcardially perfused at a rate of 2 mL/min first with PBS for 5 minutes, then with 4% PFA dissolved in PBS for 10 minutes. The brain was removed, post-fixed, and cryoprotected in 30% sucrose in PBS. Brains were cryosectioned at 20 um and slide mounted using Entellan mounting solution (Millipore). Sections were imaged at 20X objective magnification using an Axio Scan.Z1 slide scanner (Zeiss), and quantified blindly to determine the number of EB positive cells within the hilar dentate gyrus of the hippocampus.

### RT-qPCR

Total RNA was extracted from 200-250 mg of frozen mouse hippocampal tissue using TRIzol reagent (ThermoFisher Scientific #15596026), and further purified with DNA-free DNA Removal Kit (ThermoFisher Scientific AM1906). First-strand cDNA synthesis was performed from 1 μg isolated RNA template using iScript RT supermix (Bio-Rad #1708841). PCR products were amplified using a CFX96 Real-Time PCR System (Bio-Rad), and threshold cycles were detected using SsoAdvanced Universal SYBR Green Supermix (Bio-Rad #172-5271). Mean threshold cycles were normalized to 18s or GAPDH internal control, and relative gene expression levels were quantified using the 2-∆∆CT method. Primer sequences were obtained from the NCI/NIH qPrimerDepot and are listed in Table S6.

### Western Blot

Mouse or rat hippocampal tissue was homogenized and protein lysates were extracted using RIPA buffer (50 mM Tris-HCl, 150 mM NaCl, 1% NP-40, 0.5% Sodium deoxycholate, 0.1% SDS) including a protease (Calbiochem #539134) and phosphatase inhibitor cocktail (Roche PhoStop Ref: 4906845001). Protein samples were run under reducing conditions. 20 μg of protein lysate was mixed with Laemmli buffer (Bio-Rad #161-0737), containing 5% 2-mercaptoethanol (Sigma M6250), and fractionated by SDS-PAGE using the Mini-PROTEAN Tetra System and pre-cast TGX™ Gels (Bio-Rad #456-1096); Following separation, samples were transferred to a nitrocellulose membrane (0.45 μm, Bio-Rad #1620115). Membranes were blocked for 1 hr at room temperature with 5% non-fat dry milk (Apex #20-241) or 5% BSA (Biotium #22013) in TBST (10 mM Tris, 150 mM NaCl, 0.5% Tween 20, pH 8.0), and incubated overnight at 4 °C with primary antibody. Membranes were then washed 3×10 min with TBST and incubated with secondary antibodies for 1 hr at room temperature. Membranes were washed with TBST 3×10min and visualized using chemiluminescence SuperSignal West Dura Extended Substrate (ThermoFisher Scientific #34075), and Bio-Rad Chemidoc system with Bio-Rad Image Lab software (version 4.0.1). Densitometry analysis was done using Image J (NIH). The following primary and secondary antibodies were used: rabbit anti-β-Actin (1:2000, Cell Signaling #4970), rabbit anti-GAPDH (1:2000, Cell Signaling #2118), rabbit anti-TGFβR2 (1:1500, Abcam ab186838), rabbit anti-phosphorylated Smad2 (1:1000, Millipore AB3849-I), rabbit anti-phosphorylated Smad2 (1:1000, Millipore AB3849), anti-rabbit TGFβ1 (1:500, ab92486), anti-rabbit HRP (1:2000, Cell Signaling #4970).

### Osmotic pump implants

For mice, pumps were implanted as previously described*(98)*. Briefly, surgery was performed on adult male mice under isofluorane anesthesia (2%). Using a stereotaxic frame, a hole was drilled through the skull at 0.5 mm posterior, 1 mm lateral to bregma, and a cannula was placed into the right lateral cerebral ventricle, fixed with surgical glue. Cannulas (Brain infusion kit BIK 3, #0008851, Alzet, Cupertino, CA) were attached to micro-osmotic pumps (Model 2001, ALZET) filled with 200 μL of either 0.4 mM bovine serum albumin (Alb; Sigma-Aldrich) solution or artificial cerebrospinal fluid (aCSF) as previously described *(99)*, and implanted subcutaneously in the right flank. In a subset of animals, 10% of the Alb was replaced with Alexa Fluor 647 conjugated BSA (2.68 g/L; ThermoFisher Scientific A34785). Pumps infused at a rate of 1.0 μL/hr for the duration described in each experiment. For rats, 10 week-old male Wistar rats were used, and surgeries were performed the same way, using the following coordinates: −1 mm posterior and 1.5 mm lateral to bregma. For rats, albumin was used at a concentration of 0.2 mM and infused for 7 days at a rate of 10 μL/hr via larger micro-osmotic pumps (Model 2ML1, Alzet).

### ECoG

ECoG was recorded as previously reported *(49)* from 9- to 12-wk-old Wistar male rats implanted with osmotic pumps, and from young (3 months, n = 5) and old (18-22 months, n = 20) mice. In brief, under stereotaxic surgery and 2% isoflurane anesthesia, two screw electrodes were implanted in each hemisphere (rat coordinates: 4.8 mm posterior, 2.7 mm anterior, and 2.2 mm lateral; mouse coordinates: 0.5 and 3.5 mm posterior and 1 mm lateral, all relative to bregma). A wireless transmitter (Data Science International, Saint Paul, MN, US) was placed in a dorsal subcutaneous pocket, and leads connected to the screws. Connections were isolated and fixed by bone cement such that one ECoG channel was associated with each hemisphere. For rat surgeries, a cannula (−1 mm caudal, 1.5 lateral, 4 mm depth) and osmotic pump (infusing at 2.4 μl/h) were also implanted *(28)*. Animals were treated with post-operative buprenorphine (0.1 mg/Kg) and allowed to recover for at least 72 hrs prior to recording. Continuous, bichannel ECoG (sampling rate, 500Hz) was recorded wirelessly from freely roaming animals in the home cage for the duration of experiments described. Rats were recorded during the first and the fourth week following surgery. Mice were recorded for 15 days, starting 1 week after surgery, and administered IPW (20 mg/kg) on days 6-10 of recording. To detect pSWE, ECoG signals were buffered into 2 sec long epochs with an overlap of 1 sec. Fast Fourier Transform (FFT) was applied and the frequency of median power was extracted for each epoch. Thus, an event was considered as a pSWE if its frequency of median power was less than 5 Hz for 10 consecutive seconds or more. FFT was also applied for the entire recording period to analyze relative power across the frequency spectrum of 1-20 Hz.

### Animal care and transgenic mice

All animal procedures were approved by the institutional animal care committees. Animals were housed with a 12:12 light:dark cycle with food and water available ad libitum. Aldh1L1-eGFP mice were bred from STOCK Tg(Aldh1l1-EGFP)OFC789Gsat/Mmucd (identification number 011015-UCD), purchased from the Mutant Mouse Regional Resource Center. These FVB/N mice were crossed to a C57BL/6 genetic background. The resulting strain exhibits constitutive astrocytic expression of eGFP protein under the astrocytic promoter Aldh1L1.

Triple transgenic aTGFβR/KD mice were bred from strains purchased from the Jackson Laboratory to generate mice that express CreERT under the astrocytic promoter glial high affinity glutamate transporter (GLAST), with a floxed exon 4 of TGF-βR2 (tgfbr2^fl^), and a transgenic LacZ reporter gene inhibited by a floxed neomycin cassette. Tamoxifen induction thus induces activation of astrocytic CreERT resulting in a null TGFβR2 allele (tgfbr2null) and LacZ expression (R26R^−/−^). The parental strain STOCK Tg(Slc1a3-cre/ERT)1Nat/J mice were outcrossed with B6;129-Tgfbr2tm1Karl/J and B6.129S4-Gt(ROSA)26Sortm1Sor/J mice to produce males, while B6;129-Tgfbr2tm1Karl/J and B6.129S4-Gt(ROSA)26Sortm1Sor/J mice were outcrossed to produce females. The resulting GLAST-CreERT; tgfbr2^fl/+^ males were bred with tgfbr2fl/+; R26R−/− and tgfbr2^fl/fl^; R26R^−/−^ females to produce triple transgenic offspring. Subsequent generations were incrossed to produce experimental triple transgenic mice of genotypes GLAST-CreERT; tgfbr2^fl/fl^; R26R^−/−^, GLAST-CreERT; tgfbr2^fl/+^; R26R^−/−^, GLAST-CreERT; tgfbr2^fl/fl^; R26R^−/+^, and GLAST-CreERT; tgfbr2^fl/+^; R26R^−/−^. All mice were genotyped via PCR analysis of tissue biopsy samples (Table S4).

The inducible Cre/lox system was activated by 5 days of tamoxifen injection (Sigma-Aldrich, 160 mg/kg dissolved in corn oil, i.p.). Control GLAST-CreERT; tgfbr2^fl/+^ heterozygotes received the same dosage of tamoxifen and control GLAST-CreERT; tgfbr2^fl/fl^ mice received i.p. injection of corn oil vehicle at equivalent volumes. Mice were weighed daily to ensure accurate dosage.

### Seizure induction

Seizures were induced by a single injection of pentylenetetrazole (PTZ; Sigma, P6500, 85 mg/kg s.c.). Video recordings were taken for 30 minutes following induction, which were then scored by a blind observer, using a modified Racine scale to quantify progression of seizure severity, as follows: 0 – No Seizures; 1 – Immobility; 2 – Straub’s tail; 3 – Forelimb Clonus; 4 – Generalized Clonus; 5 – Bouncing Seizure; 6 – Status Epilepticus. Latency to first appearance of each Racine stage was quantified, as was latency to mortality (when occurring prior to the 30 min endpoint).

### Behavior assessments

Spatial working memory was tested by spontaneous alternation in a T-maze constructed of black plastic, with stem (50 x 16 cm) and two arms (50 x 10 cm). A vertical divider was placed at the midline of the stem exit, thus creating two entryways leading to either the right or left arms. Naïve mice were placed at the beginning of the stem and allowed to freely roam until opting to enter either the right or left arm. Upon crossing the threshold of the chosen arm, a door was lowered, and the mouse was contained to the chosen arm for 30 sec. Then, the mouse was returned to the stem and the door raised, allowing the next choice trial to commence immediately. After 10 sequential trials, the mouse was returned to the home cage. “Correct” alternation choices were scored when the mouse chose the arm opposite of that chosen in the previous trial, and percent correct was calculated as (total correct choices) / (total number of completed trials). In the event that a mouse did not leave the stem within 60 sec, it was removed from the apparatus for 30 sec, and then reset in the stem to start a new trial. These “incomplete” trials were not counted in scoring. Spatial memory in young mice was assessed in the Morris Water Maze (MWM) task *(100)*. Because of excess colony availability, heterozygous control mice from the transgenic colony (TGFβR2 (fl/+) genotype), which retain normal TGFβ signaling, were used for the MWM study cohort. For each trial, mice were placed at randomized starting locations in a MWM pool filled with opaque water (colored with non-toxic white acrylic paint), with visual cues placed on the pool walls. Mice were allowed to swim freely until locating a hidden platform under the surface of the water (or guided to the platform after 40 seconds of failed searching), and then left on the platform for 10 seconds prior to starting a new trial. Spatial learning was quantified by measuring latency to reach the platform, averaged from four trials per day, with training over 9 consecutive days of learning. Object memory was tested in the novel object task, using a 3-day protocol consisting of a 10 min trial on each day, recorded by overhead video. On day 1 mice were habituated to a square testing chamber, constructed of white plastic (50 x 50 cm), with two unfamiliar objects placed inside. On day 2, the previous objects were removed and 3 new objects were placed in a “L” configuration, equidistant from each other and the walls. On day 3, one of the objects was removed and replaced by a new object, thus leaving 2 familiar objects and 1 novel object. A blind observer quantified duration of time spent investigating each object (scoring criteria were mouse nose oriented towards object at a distance of 1 cm or less), and percentage of time at the novel object was quantified as (duration investigating novel object) / (total duration investigating all three objects). The set objects were chosen from common items such as lab bottles, pipette boxes, cups (placed open-side down), etc., that were of similar dimensions but varied in shape, color, and material. The sequence and position of objects used across trials was identical for all mice.

### IPW pharmacokinetics

Brain concentration measurement of IPW-5371 was performed by Jubilant Biosys. The experiment was approved by the Jubilant Biosys Institutional Animal Ethics Committee, Bangalore, India (IAEC/JDC/2015/72) and were in accordance with the Committee for the Purpose of Control and Supervision of Experiments on Animals (CPCSEA), Ministry of Social Justice and Environment, Government of India. Twelve Balb/C mice (age 6-7 weeks) were procured from Bioneeds, Bangalore, India. Animals were housed in Jubilant Biosys animal care facility in a temperature and humidity controlled room with a 12:12 h light:dark cycles, had free access to food (Provimin, India) and water for one week before experimental use. Following ~4 h fasting (during fasting period animals had free access to water) mice received IPW-5371 orally at a dose of 20 mg/kg (dissolved in 0.5% methylcellulose and saline). Mice were euthanized under isofluorane and the brain was removed and weighed. Brain tissue homogenates were prepared with 10% tetrahydrofuran in acetonitrile (tissue was homogenated with 4 mL/g of 10% tetradhydrofuran in acetonitrile containing IS (100 ng/mL)). Subsequently, 250 μL of brain homogenate was centrifuged at 14,000 rpm for 5 min at 10 °C. An aliquot of 10 μL was injected onto an API 4000 LC-MS/MS system for analysis.

## Acknowledgements

This research was supported by NSF GRFP fellowships (V.V.S. and A.R.F.); Siebel Fellowship (V.V.S); NIH NRSA fellowship F31AG054147 (A.R.F.); NIH grants R01NS066005 and R56NS066005, a Bakar Foundation Fellowship, the Archer Foundation Award (D.K.); the European Union’s Seventh Framework Program (FP7/2007-2013, grant agreement 602102, EPITARGET), the Israel Science Foundation (717/15) (A.F.); NIH grant R01AG042679 (A.D.); and the Binational Israel-USA Science Foundation (D.K. and A.F.). We thank Kelley Patten, Ami Citri, Hermona Soreq, and Inbal Goshen for edits on the manuscript. We thank Su Lee for graphic design of the schematic diagram.

