## Supplemental Information for "Blood-brain barrier dysfunction in aging induces hyper-activation of TGF-beta signaling and chronic yet reversible neural dysfunction"

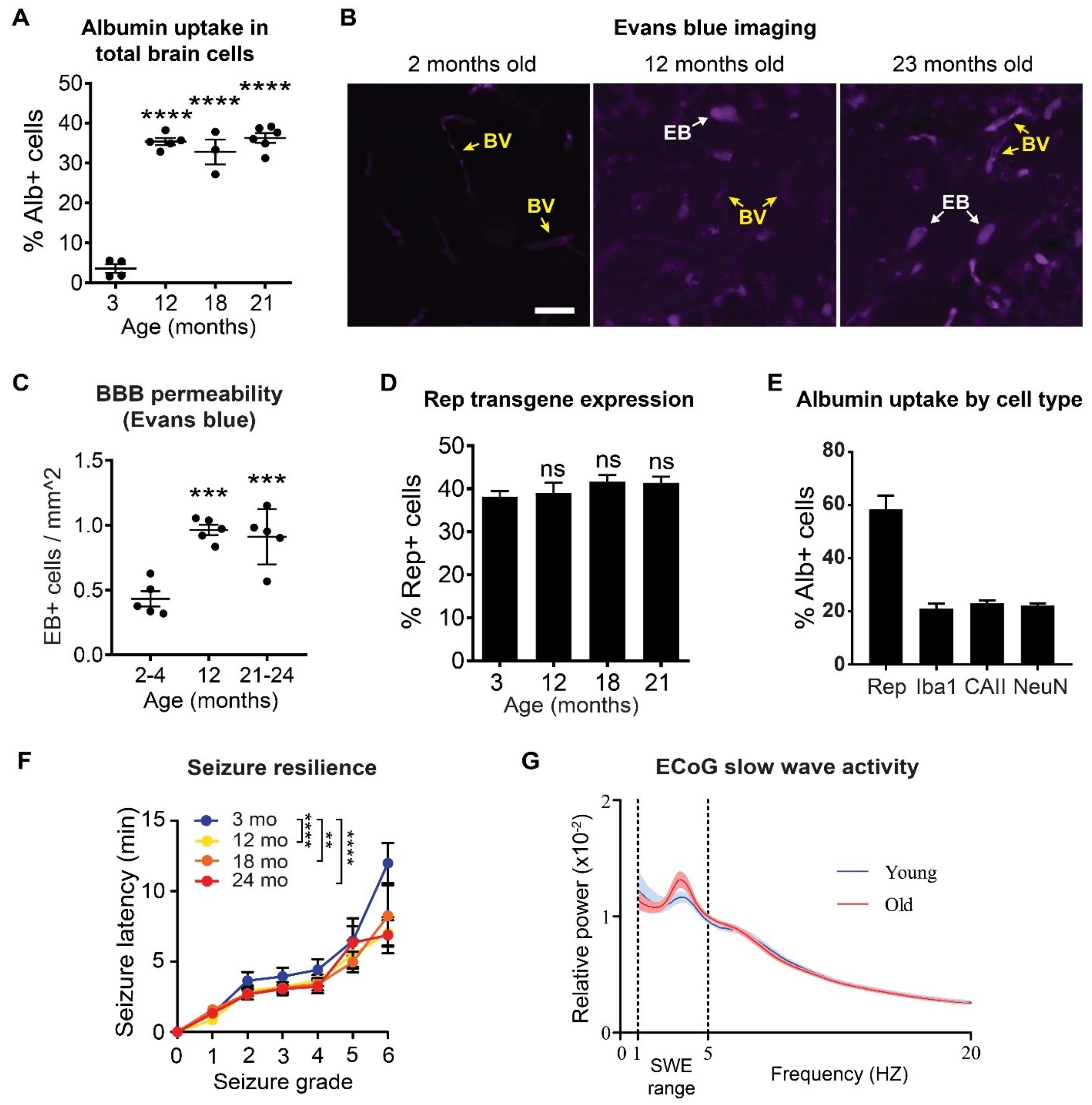


**Figure S1.** (A) Percentage of cells (DAPI+) in the mouse hippocampus that colocalize with albumin at given timepoints across the mouse lifespan; albumin extravasation and localization to brain cells increases significantly with age (ANOVA, $p < 0.0001$, with Bonferroni posttest). (B-C) BBB breakdown in aging mice was confirmed by the Evans blue (EB) assay. 30 min after i.v. EB injection, the EB tracer was mostly contained within blood vessels in young mice (2-4 month old), while middle aged (12 month old) and old (21-24 month old) mice showed significantly elevated extravascular EB tracer localized to cells (ANOVA, n = 5, $p < 0.0002$, with Dunnett's posttest). Representative images from mouse hilar hippocampal dentate gyrus show EB (purple) localized to blood vessels (BV, yellow arrows) and extravascular EB (EB, white arrows) localized to cells. Scale bar = 20 µM. (D) Percent of total cells expressing fluorescent reporter (Rep) in the hippocampus of Rep-Aldh1L1 mice. The number of labeled astrocytes did not change across the lifespan. (E) Co-staining for cell-specific markers was used to estimate the percent of albumin taken up by different cell types in the aged mouse hippocampus, as a percent of all albumin positive cells. Albumin was predominantly taken up by astrocytes (Rep), and was found at lower levels in microglia (Iba1), oligodendrocytes (CAII), and neurons (NeuN). (F) A modified Racine scale, measuring seizure severity on a scale of 1-6, was used to score seizure vulnerability following injection of PTZ. Aged mice showed a faster Racine progression (onset of first seizure at each stage of severity; 2-way ANOVA, main effect of seizure progression $p < 0.0001$; main effect of age $p = 0.005$, n = 13 (3 mo), 10 (12 mo), 8 (18 mo and 24 mo)). For each group, linear regression was fit to the Racine progression, and the slope of the best fit line was used as a measure of seizure vulnerability in Fig. 1F. (G) Spectral analysis of ECoG showed an increase in slow-wave power in aged mice compared to young. This slow-wave activity was examined in greater detail by quantifying discrete pSWEs (see Fig. 1H-I).

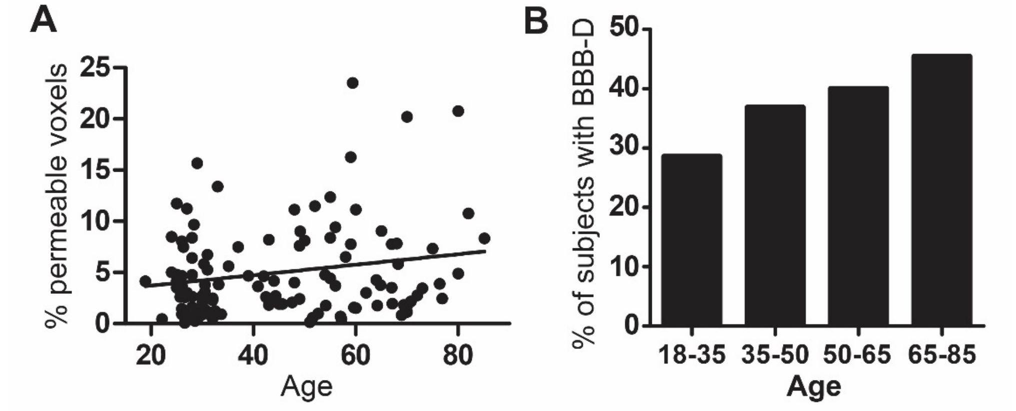


**Figure S2.** (A) Linear regression was used to model the relationship between age and BBB permeability (% of total brain volume with permeability values above the 95th percentile of the averaged young healthy subjects), showing a significant increase with age (n = 113, p=0.0315). (B) Descriptive statistics estimating the percent of individuals affected by BBB-D (defined as suprathreshold permeability in more than 5% all brain voxels) across age groups.

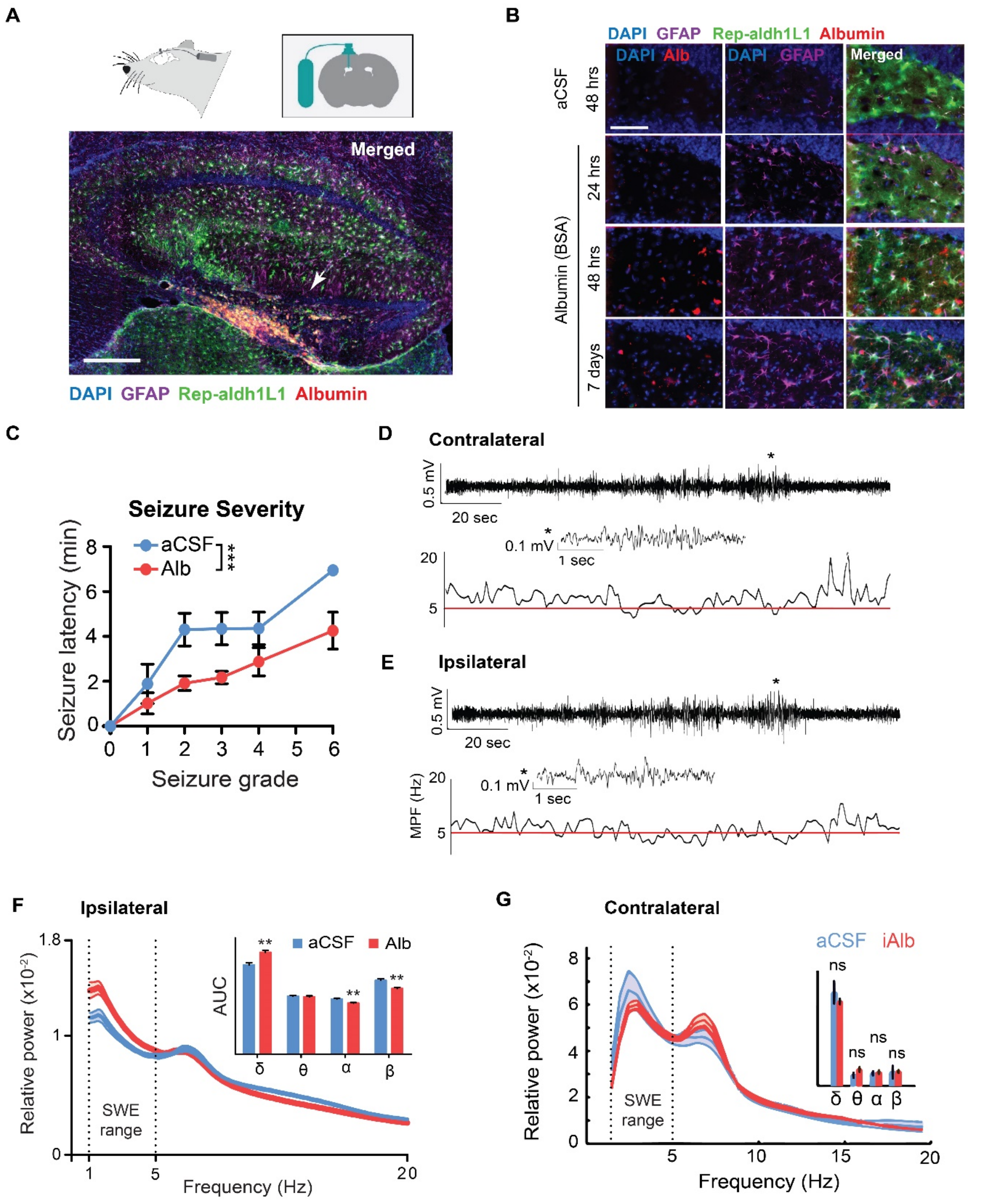


**Figure S3.** (A) (Top) Diagrams showing mouse icv pump implants, with cannulas placed to infuse albumin into the brain ventricle; (Bottom) Immunofluorescent image of ipsilateral hemisphere, showing hippocampus (coronal slice). The infused albumin (fluorescent-tagged, in red) diffuses into the ventricle and accumulates in the dentate gyrus (arrowhead indicates the granule cell layer). Scale bar = 700 µm. (B) Representative images from the dentate gyrus show albumin uptake in astrocytes at 24 hours, 48 hours, and 7 days after pump implant (with aCSF control shown at 48 hours; scale bar

= 50 μm). (C) iAlb mice showed a faster progression through the Racine scale compared to aCSF controls (2-way ANOVA, main effect of seizure progression $p < 0.0001$; main effect of treatment $p = 0.0001$, $n = 4$). Best fit lines through the Racine progressions were used to measure seizure vulnerability in Fig. 2B. (D-E) Representative ECoG traces from ipsilateral and contralateral hemisphere, showing discrete pSWEs within a 10 second window (marked with *). (F) Rats showed an increase in slow-wave power in the ipsilateral hemisphere receiving iAlb infusion, whereas no shift in the spectrum of ECoG activity was observed in recordings from (G) contralateral hemisphere.

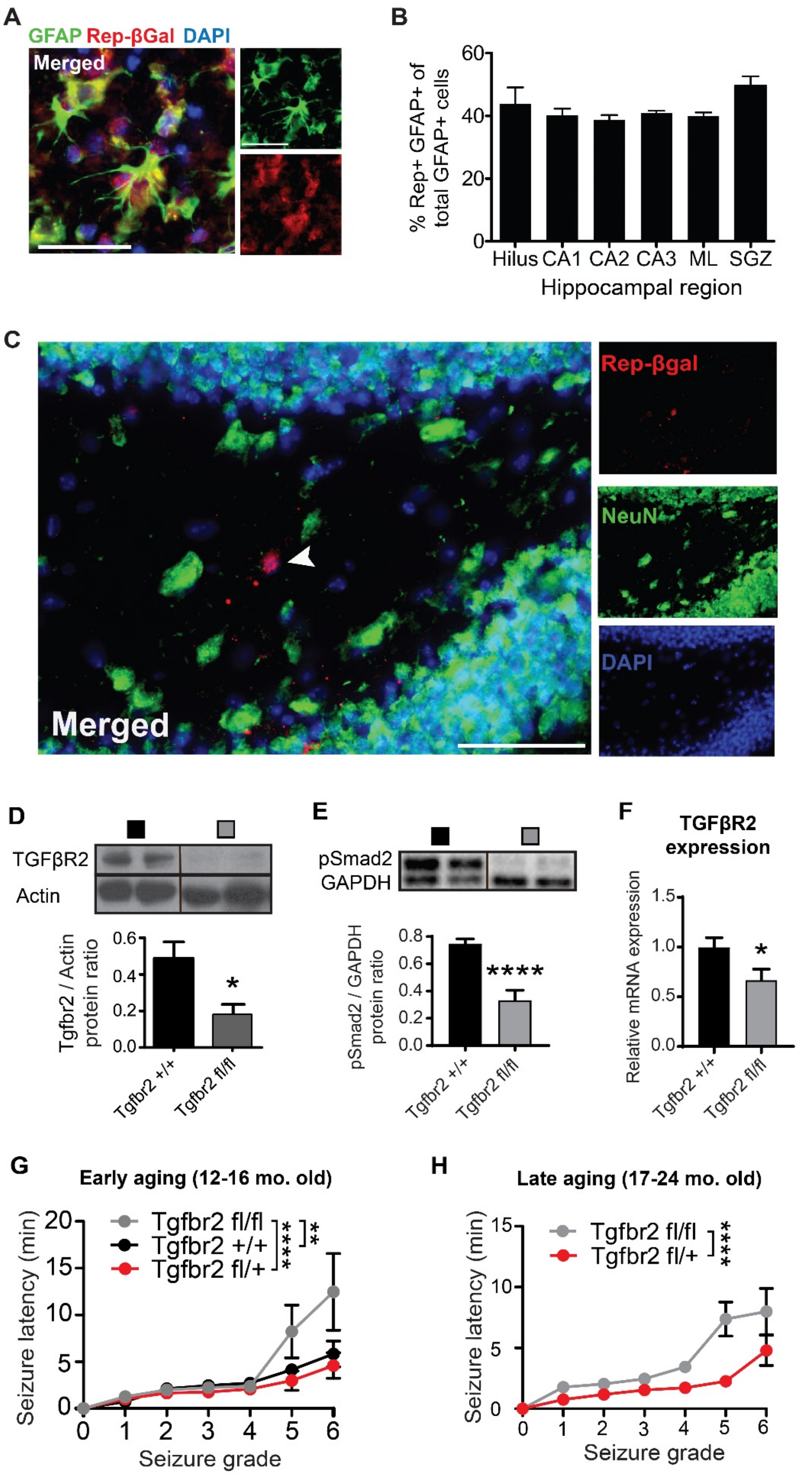

A
GFAP Rep-βGal DAPI
Merged
B
% Rep+ GFAP+ of total GFAP+ cells
Hilus CA1 CA2 CA3 ML SGZ
Hippocampal region
C
Rep-βgal
NeuN
DAPI
Merged
D
TGFβR2
Actin
Tgfbr2 / Actin protein ratio
Tgfbr2 +/+
Tgfbr2 fl/fl
E
pSmad2
GAPDH
pSmad2 / GAPDH protein ratio
F
TGFβR2 expression
Relative mRNA expression
G
Early aging (12-16 mo. old)
Tgfbr2 fl/fl
Tgfbr2 +/+
Tgfbr2 fl/+
Seizure latency (min)
Seizure grade
H
Late aging (17-24 mo. old)

**Figure S4.** (A) Representative immunofluorescent image from mouse hippocampus, showing reporter expression (Rep-βGal, red) in astrocytes (GFAP, green) following induction of the Cre recombinase system to knock down (KD) aTGFβR. Scale Bar = 30 μm. (B) Reporter expression was present in approximately 40% of hippocampal astrocytes following induction of KD, and was consistent throughout all hippocampal subregions (CA: cornu ammonis; ML: molecular layer; SGZ: subgranular zone). (C) No significant reporter expression was observed in mature neurons (NeuN), indicating that hippocampal neural stem cells expressing GLAST did not generate an appreciable lineage of recombinant TGFβR KD neurons. Representative image of reporter expression in mouse hippocampus stained with NeuN to assess neural expression, scale bar = 50 μm. (D) Western blot with densitometry confirmed significant reduction in levels of TGFβR2 in the hippocampus following tamoxifen induction in homozygous KD mice (aTGFβR2 fl/fl) compared to controls (aTGFβR2 +/+) (t-test, p=0.045, n=3). (E) WB from 17-24 month aged mouse hippocampus showing reduced pSmad2 following KD (t-test, $p < 0.0001$, n = 9 $Tgfbr2^{+/+}$, 7 $Tgfbr2^{fl/fl}$). (F) Quantitative real-time PCR from aged mouse hippocampus showing decreased expression of TGFβR2 following KD (t-test, p = 0.036, n = 10 $Tgfbr2^{+/+}$, 8 $Tgfbr2^{fl/fl}$). (G-H) aTGFβ KD significally reduced progression through the Racine scale in early and late aged mice (2-way ANOVA; for early aged group, main effect of seizure progression $p < 0.0001$; main effect of genotype p = 0.0053; interaction p = 0.0214, with Bonferroni posttest, n = 5 $Tgfbr2^{+/+}$, 6 $Tgfbr2^{fl/+}$ and $Tgfbr2^{fl/fl}$; for late aged group, main effect of seizure progression $p < 0.0001$; main effect of genotype $p < 0.0001$; interaction p = 0.026). Best fit lines of Racine progression were used to measure seizure vulnerability in Fig. 3C and D.

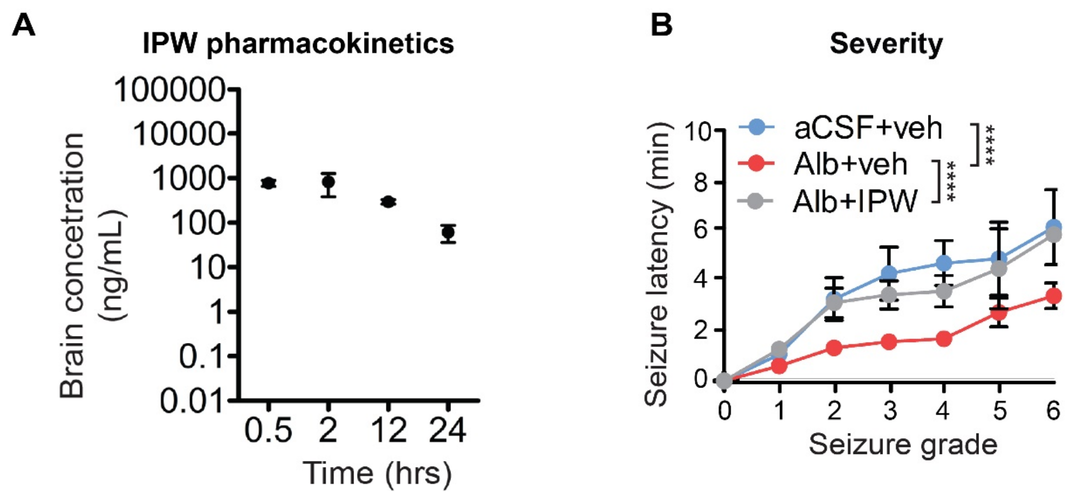


**Figure S5.** (A) PK measures of IPW concentration in brain lysates from n = 3 mice at timepoints spanning 24 hours after a single dose (30 mg/kg, po). At 24 hours, brain IPW concentration remains above the $IC_{50}$ of 75 nM. (B) IPW treatment significantly slowed progression through the Racine seizure scale in iAlb mice (2-way ANOVA, main effect of seizure progression $p < 0.0001$; main effect of treatment p = 0.0001, n = 4). Best fit lines of Racine progression were used to measure seizure vulnerability in Fig. 5C.

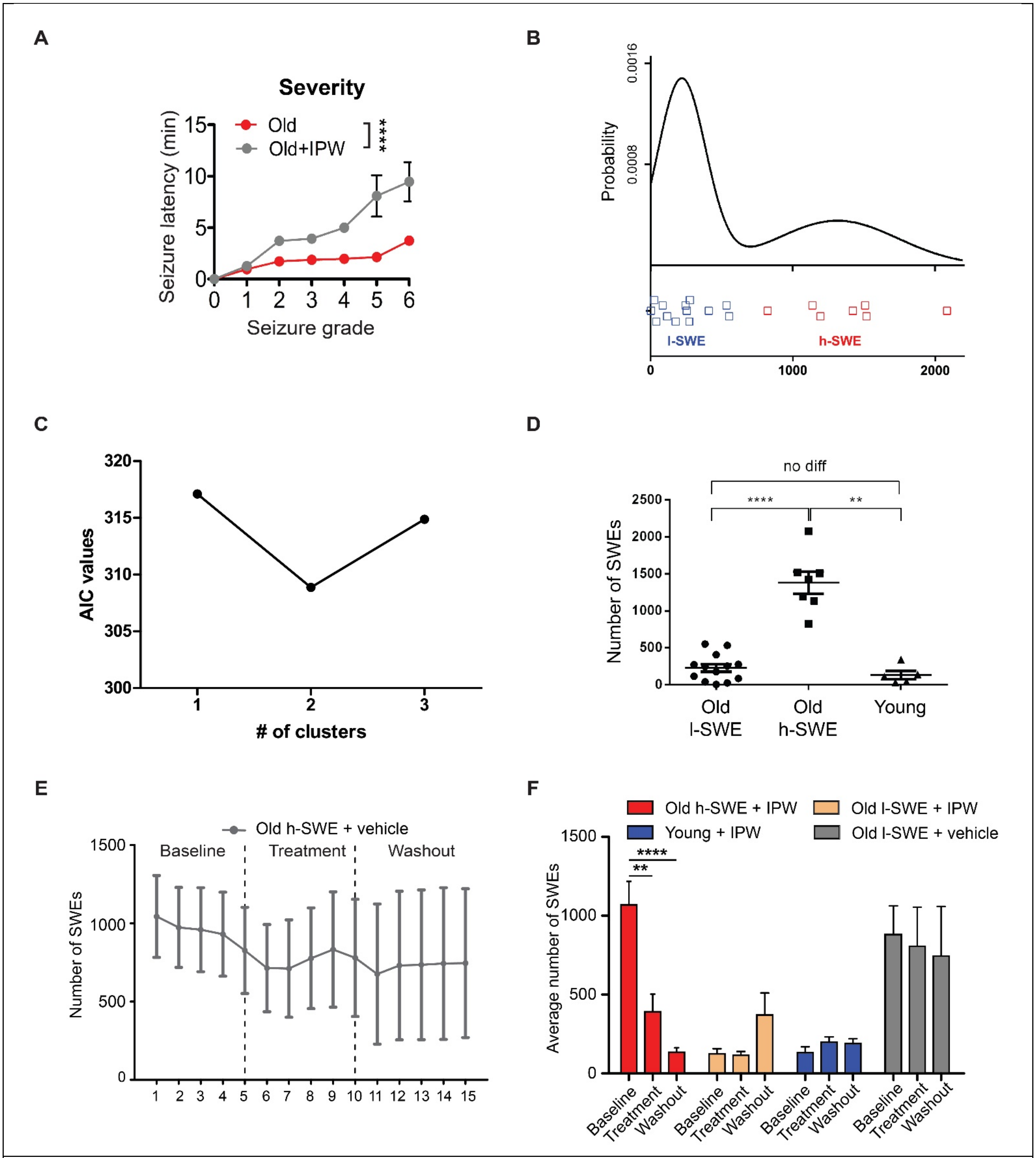


**Figure S6.** (A) 7 days of IPW treatment reversed seizure vulnerability in aged mice, reducing progression through the Racine scale (2-way ANOVA, main effect of seizure progression $p < 0.0001$; main effect of drug treatment $p < 0.0001$; interaction $p = 0.0001$, with Bonferroni posttest, $n = 9$). Best fit lines of Racine progression were used to measure seizure vulnerability in Fig. 6D. (B) GMM clustering was performed to segregate the old mice into l-SWE (asymptomatic) and h-SWE (symptomatic) subgroups. (C) Akaike information criterion (AIC) was used to compare the goodness-of-fit of GMM clustering models with 1, 2, or 3 clusters. Lower AIC values represent better goodness-of-fit, indicating the old mice are best described by the two cluster model (low and high subgroups) rather than as one continuous group. (D) The old h-SWE group had significantly more pSWEs than the old l-SWE group or young (ANOVA). (E) Mice treated with injection of vehicle (rather than IPW) showed no change in number of pSWEs. (F) Average number of pSWEs for all groups during the final two days of each treatment period (Baseline prior to injection, treatment with daily injection of IPW or vehicle control, and washout with cessation of injections). Treatment significantly reduces pSWEs in the h-SWE group, whereas no changes in pSWEs were observed in any other group.

**Table S1. Age and gender of human subjects who received DCE-MRI scans**

| Age | Gender | MRI abnormalities | Diagnosis* |
|---|---|---|---|
| 18.91 | M | None | None |
| 22.17 | F | None | None |
| 24 | M | None | None |
| 25 | F | None | None |
| 25 | M | None | None |
| 25 | M | None | None |
| 25.71 | M | None | None |
| 26 | M | None | None |
| 26 | M | None | None |
| 26 | M | None | None |
| 26 | M | None | None |
| 26.12 | F | None | None |
| 26.6 | F | None | None |
| 26.67 | F | None | None |
| 27 | M | None | None |
| 27.24 | F | None | None |
| 28 | M | None | None |
| 28 | M | None | None |
| 28 | M | None | None |
| 28 | M | None | None |
| 28.59 | M | None | None |
| 29.46 | M | None | None |
| 30 | M | None | None |
| 30 | F | None | None |
| 30 | M | None | None |
| 30 | M | None | None |
| 30 | M | None | None |
| 30.37 | F | None | None |
| 30.53 | F | None | None |
| 30.56 | M | None | None |

| | | | |
|---|---|---|---|
| 30.69 | M | None | None |
| 31 | M | None | None |
| 32 | M | None | None |
| 32 | M | None | None |
| 32.06 | M | None | None |
| 32.55 | F | None | None |
| 33.24 | F | None | None |
| 33.72 | M | None | None |
| 39 | M | None | None |
| 40.91 | M | None | None |
| 42 | M | None | None |
| 42.46 | M | None | None |
| 43 | F | None | None |
| 44 | F | None | None |
| 44.33 | M | None | None |
| 45.63 | F | None | None |
| 47.62 | F | None | None |
| 48 | F | None | None |
| 49 | F | None | None |
| 49 | F | None | None |
| 51 | M | None | None |
| 51.58 | F | None | None |
| 52.68 | F | None | None |
| 54 | F | None | None |
| 54.16 | F | None | None |
| 55 | F | None | None |
| 56 | M | None | None |
| 57 | F | None | None |
| 57.22 | F | None | None |
| 59.63 | F | None | None |
| 60 | F | None | None |
| 62 | F | None | None |
| 64 | F | None | None |
| 64.22 | F | None | None |
| 64.85 | F | None | None |
| 67 | M | None | None |
| 67 | M | None | None |
| 67.18 | M | None | None |
| 68.3 | M | None | None |
| 68.97 | F | None | None |
| 69.41 | F | None | None |
| 70 | F | None | None |
| 70.18 | F | None | None |
| 70.81 | M | None | None |

| | | | |
|---|---|---|---|
| 72 | F | None | None |
| 73.02 | M | None | None |
| 76.4 | F | None | None |
| 76.89 | F | None | None |
| 80 | M | None | None |
| 24 | M | None | None |
| 25 | M | None | None |
| 26 | M | None | None |
| 26.36 | M | None | None |
| 27 | M | None | None |
| 28 | M | None | None |
| 28 | M | None | None |
| 28.39 | M | None | None |
| 29 | M | None | None |
| 31 | M | None | None |
| 33 | M | None | None |
| 35.13 | F | None | None |
| 37 | F | None | None |
| 41 | M | None | None |
| 43 | F | None | None |
| 48 | F | None | None |
| 49.16 | F | None | None |
| 50 | F | None | None |
| 52.02 | F | None | None |
| 55 | M | None | None |
| 55 | F | None | None |
| 56 | M | None | None |
| 58 | F | None | None |
| 59 | F | None | None |
| 59 | M | None | None |
| 59.4 | M | None | None |
| 60 | M | None | None |
| 65 | M | None | None |
| 68 | F | None | None |
| 70 | M | None | None |
| 75 | F | None | None |
| 80 | M | None | None |
| 82 | M | None | None |
| 85 | M | None | None |
| 85.17 | M | None | None |

*Patients were screened for confounding neurological conditions, including previous diagnosis of stroke, head injury, and cognitive impairment.

### Table S2. Gender and age of subjects providing post-mortem brain tissue

| Gender | Age | Diagnosis* |
|---|---|---|
| m | 26 | None |
| m | 32 | None |
| f | 36 | None |
| m | 61 | None |
| f | 63 | None |
| m | 68 | None |
| f | 69 | None |
| m | 70 | None |
| m | 71 | None |
| m | 74 | None |
| f | 75 | None |
| m | 77 | None |
| m | 78 | None |

*Subjects were screened for any confounding neurological condition, including stroke, head injury, and dementia/neurodegenerative disease.

### Table S3. PCR primers used to perform genotyping

| Primer | Sequence | Amplicon Size (bp) |
|---|---|---|
| Tg(Slc1a3-cre/ERT)1Nat fwd | 5' ACA ATC TGG CCT GCT ACC AAA GC 3' | Transgene: ~600 |
| Tg(Slc1a3-cre/ERT)1Nat rev | 5' CCA GTG AAA CAG CAT TGC TGT C 3' | Internal positive control: 200 |
| Tg(Slc1a3-cre/ERT)1Nat IPC 1 | 5' CAA ATG TTG CTT GTC TGG TG 3' | |
| Tg(Slc1a3-cre/ERT)1Nat IPC 2 | 5' GTC AGT CGA GTG CAC AGT TT 3' | |
| Tgfbr2$^{tm1Karl}$ fwd | 5' TAT GGA CTG GCT GCT TTT GTA TTC 3' | Mutant: 575 |
| Tgfbr2$^{tm1Karl}$ rev | 5' TGG GGA TAG AGG TAG AAA GAC ATA 3' | Heterozygote: 415 and 575 |
| | | Wild type: 415 |
| Gt(ROSA)26Sor$^{tm1Sor}$ common | 5' GCG AAG AGT TTG TCC TCA ACC 3' | Mutant: 340 |
| Gt(ROSA)26Sor$^{tm1Sor}$ mutant rev | 5' AAA GTC GCT CTG AGT TGT TAT 3' | Heterozygote: 340 and ~650 |
| Gt(ROSA)26Sor$^{tm1Sor}$ wild type rev | 5' GGA GCG GGA GAA ATG GAT ATG 3' | Wild Type: ~650 |
| eGFP fwd | 5' CCT ACG GCG TGC AGT GCT TCA GC 3' | eGFP: ~300 |
| eGFP rev | 5' CGG CGA GCT GCA CGC TGC GTC CTC 3' | |

### Table S4. Components of PCR Reactions

| Allele | Reaction Component | Volume per Reaction (μL) |
|---|---|---|
| Tg(Slc1a3-cre/ERT)1Nat | | |

| | | |
|---|---|---|
| | Forward Primer | 4 |
| | Reverse Primer | 4 |
| | IPC 1 Primer | 2 |
| | IPC 2 Primer | 2 |
| | Apex 2X Taq RED Master Mix | 12.5 |
| | DNA Template | 0.5 |
| | Total | 25 |
| Tgfbr2tm1Karl | | |
| | Forward Primer | 2.5 |
| | Reverse Primer | 2.5 |
| | Apex 2X Taq RED Master Mix | 12.5 |
| | Water | 7 |
| | DNA Template | 0.5 |
| | Total | 25 |
| Gt(ROSA)26Sortm1Sor | | |
| | Common Primer | 2.5 |
| | Mutant Reverse Primer | 2.5 |
| | Wild Type Reverse Primer | 2.5 |
| | Apex 2X Taq RED Master Mix | 12.5 |
| | Water | 4 |
| | DNA Template | 0.5 |
| | Total | 25 |
| eGFP | | |
| | Forward Primer | 2.5 |
| | Reverse Primer | 2.5 |
| | Apex 2X Taq RED Master Mix | 12.5 |
| | Water | 7 |
| | DNA Template | 0.5 |
| | Total | 25 |

### Table S5. PCR Thermocycling Conditions

| Allele | Cycling Step # | Temp (°C) | Time | Note |
|---|---|---|---|---|
| Tg(Slc1a3-cre/ERT)1Nat | | | | |
| | 1 | 94 | 3:00 | - |
| | 2 | 94 | 0:30 | - |
| | 3 | 60 | 0:30 | - |
| | 4 | 72 | 0:30 | - |
| | 5 | - | - | Go to 2, 35X |
| | 6 | 72 | 2:00 | - |
| | 7 | 12 | 0:00 | infinite hold |
| Tgfbr2tm1Karl | | | | |

| | | | | |
|---|---|---|---|---|
| | 1 | 94 | 3:00 | - |
| | 2 | 94 | 0:30 | - |
| | 3 | 62 | 0:30 | - |
| | 4 | 72 | 0:30 | - |
| | 5 | - | - | Go to 2, 35X |
| | 6 | 72 | 2:00 | - |
| | 7 | 10 | 0:00 | infinite hold |
| Gt(ROSA)26Sortm1Sor | | | | |
| | 1 | 98 | 3:00 | - |
| | 2 | 98 | 0:30 | - |
| | 3 | 65 | 1:00 | - |
| | 4 | 72 | 1:00 | - |
| | 5 | - | - | Go to 2, 35X |
| | 6 | 72 | 2:00 | |
| | 7 | 10 | 0:00 | infinite hold |
| eGFP | | | | |
| | 1 | 94 | 3:00 | - |
| | 2 | 94 | 0:30 | - |
| | 3 | 60 | 0:45 | - |
| | 4 | 72 | 0:45 | - |
| | 5 | - | - | Go to 2, 35X |
| | 6 | 72 | 10:00 | - |
| | 7 | 11 | 0:00 | infinite hold |

### Table S6. Primers for qPCR

| | |
|---|---|
| Tgfbr2-F | CTGGCCATGACATCACTGTT |
| Tgfbr2-R | GTCGGATGTGGAAATGGAAG |
